# Detecting and Correcting False Transients in Calcium Imaging

**DOI:** 10.1101/473470

**Authors:** Jeff L. Gauthier, Sue Ann Koay, Edward H. Nieh, David W. Tank, Jonathan W. Pillow, Adam S. Charles

## Abstract

Population recordings of calcium activity are a major source of insight into neural function. Large dataset sizes often require automated methods, but automation can introduce errors that are difficult to detect. Here we show that automatic time course estimation can sometimes lead to significant misattribution errors, in which fluorescence is ascribed to the wrong cell. Misattribution arises when the shapes of overlapping cells are imperfectly defined, or when entire cells or processes are not identified, and misattribution can even be produced by methods specifically designed to handle overlap. To diagnose this problem, we develop a transient-by-transient metric and a visualization tool that allow users to quickly assess the degree of misattribution in large populations. To filter out misattribution, we also design a robust estimator that explicitly accounts for contaminating signals in a generative model. Our methods can be combined with essentially any cell finding technique, empowering users to diagnose and correct at large scale a problem that has the potential to significantly alter scientific conclusions.

## 1 Introduction

Many recent advances in understanding nervous system function have relied on large-scale population recordings at cellular resolution. A principal method in this family of techniques is the expression of genetically-encoded calcium indicators in a set of neurons [2, 26, 1], followed by one- or two-photon imaging of hundreds or thousands of cells or processes [5, 8, 28]. As advances in microscopy and indicator expression [4, 20, 2, 1] have allowed for larger, longer, and more numerous recording sessions, it has become essentially impossible for users to manually identify every active cell in every movie frame. Thus automated methods have become indispensable for analyzing large datasets. Automation, however, creates a layer of separation between users and raw data that has the potential to mask errors.

The principal task when analyzing calcium imaging movies is to assign a time course of fluorescence changes to each source of activity, with particular emphasis on the transient increases in brightness following spiking events [2]. Active sources include cell bodies, dendrites, axons, and neuropil, a mesh of small processes in which individual structures can not be distinguished. A typical workflow involves first identifying the spatial extent, or “profile”, of each source using either manual or automated methods [7, 25, 11, 28, 24] (see glossary for list of naming conventions). Inferring the time courses of neural activity from raw fluorescence measurements, however, is always performed automatically, either by averaging pixels in each region of interest [7, 25], or employing more sophisticated algorithms [28, 11].

It has long been recognized that overlapping sources can contaminate one another’s time courses, and that unwanted mixing of signals has the potential to distort scientific conclusions [6, 19, 18, 12]. Most methods for source identification are designed to prevent such contamination, which raises the question of how to assess their effectiveness. In the case of neuropil mixing with cell bodies, the neuropil time course around each cell is readily estimated by drawing a small annulus, and de-contamination can be verified by showing that this neuropil signal is no longer present in the cell time course [28, 16, 17, 29]. Here, however, we are concerned with identifying contamination from non-neuropil sources, such as when a dendrite overlaps a cell body, or two cell bodies overlap.

In the absence of “ground truth”, contamination can be difficult to diagnose. Typical methods of validation entail quantifying the explained variance, cell-wise correlations to the average movie [11], or statistical tests on the temporal residual [21]. Even when simulations provide ground-truth data [25], or are performed on a cell-wise basis [11], evaluation metrics are global and do not examine individual transients (though single spikes have been considered in the special case of simultaneous electrical and optical recording of Purkinje cell complex spikes [25]). Thus available methods lack a generalizable framework for verifying that each transient was produced by its assigned source.

Here, we investigate contamination in two-photon imaging, how it might affect scientific conclusions, and develop tools to explore, quantify, and remedy such errors. As a test case, we examine a brain region where contamination is particularly likely to occur: the CA1 region of the hippocampus. CA1 cell bodies are densely packed, with essentially no gap between neighboring cells, and are intermingled with thick apical dendrites [6], creating many opportunities for significant overlap between sources. By performing a close comparison of movie frames and identified source profiles, we will develop automated measures to assess the level of contamination in source time courses. In addition, we will present a new time course estimation algorithm to explicitly model and filter out activity assigned in error.

## 2 Results

### 2.1 False transients occur and can impact scientific conclusions

We begin by exploring how inferred activity time courses relate to the original motion-corrected fluorescence movie. Our purpose is not to evaluate a particular cell finding algorithm or consider how it might be improved. Instead, we aim to develop a suite of tools that allow users to perform their own evaluation, one that can be tailored to meet the needs of each application. As a test case, we examine a movie recorded in dorsal CA1 [10]. Cell finding was performed using a modern algorithm, Constrained Non-negative Matrix Factorization (CNMF) [31], to ensure the results were representative of current methods. Time courses were estimated by fitting source profiles to each frame independently with least squares, since additive Gaussian noise (which motivates least squares) is a common basis for more sophisticated modeling, e.g. auto-regressive temporal processes [31].

The initial diagnosis focuses on a small patch in which 52 sources were identified (Fig. 1A). Upon simple inspection, source profiles seem to match the cells in the average image, suggesting that cell finding was generally accurate in identifying source shapes. Similarly, the time course of each source matches the expected statistics of calcium dynamics in this region: a low baseline level of fluorescence punctuated by transient increases, a collection of which are shown for one source (Fig. 1B, see Methods for transient definition).

**Figure 1:**
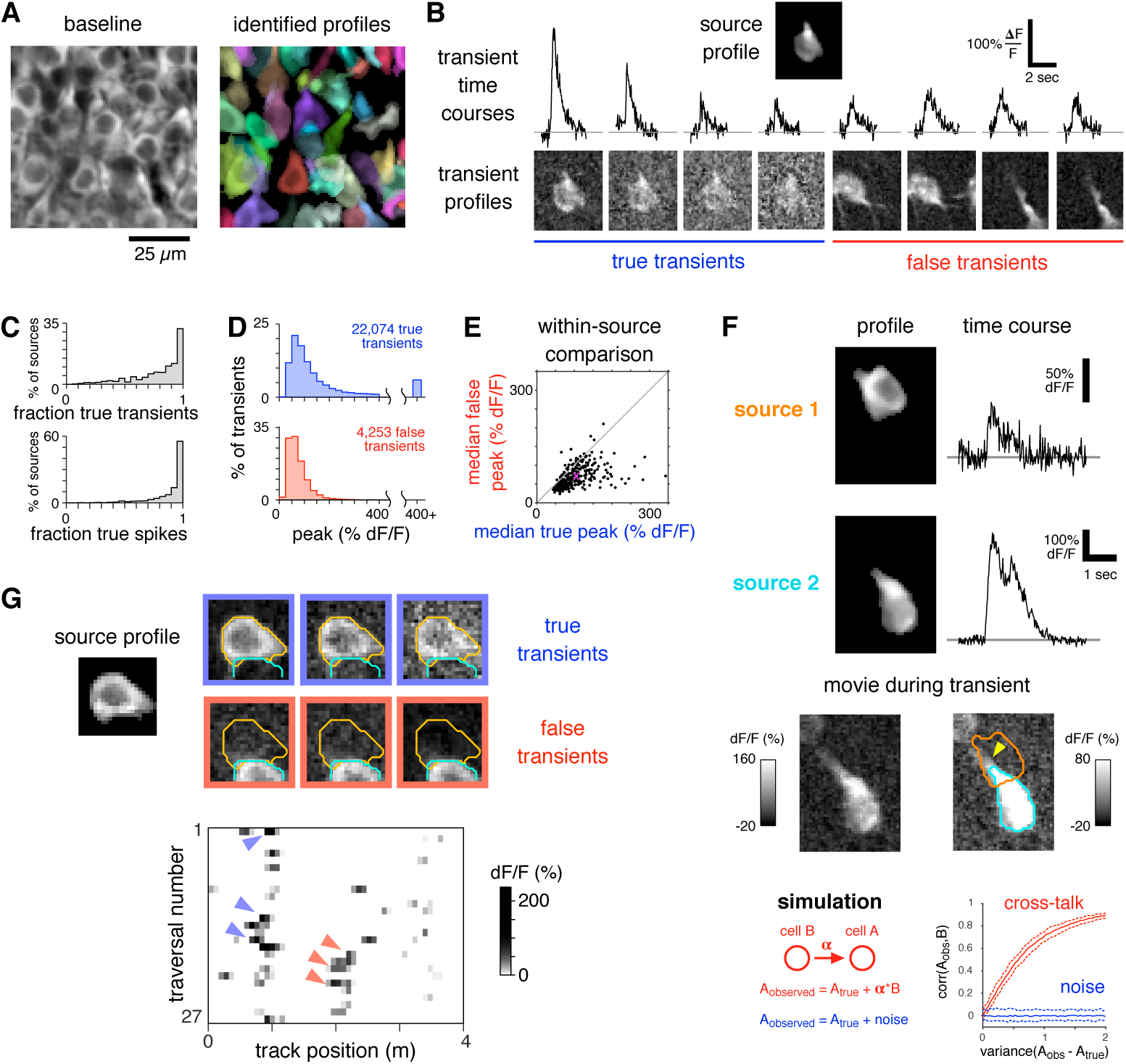
False transients occur and can interfere with scientific results. A: Left: Mean fluorescence image from a subset of an *in vivo* two-photon imaging movie showing mouse CA1 pyramidal cells genetically expressing GCaMP3. Right: Spatial profiles of the 52 sources identified by CNMF in this movie, each a different color. B: Time courses (top row) of eight transients assigned to one source (top inset image) and their respective transient profiles (bottom row). Classification as true or false was performed manually by a human expert. C: Occurrence of true activity in a set of 1,325 simultaneously-recorded sources. Top shows fraction of transients which were classified as true (only for sources with at least 1 classified transient). Bottom shows equivalent data for spikes that were inferred from classified transients (only for sources with at least 1,000 spikes inferred from classified transients). D: Peak amplitudes of true and false transients for the same sources as in C. E: Median peak amplitude of true and false transients for each source, only shown for sources with at least 5 true and 5 false transients. Purple “X” indicates mean over sources. F: Top panels: Two source profiles and their respective time courses during a brief period of the movie. Middle panels: transient profile for source 2 (left), and the same image shown at increased contrast with overlaid outlines of source profiles (right). Arrowhead indicates part of source 2 fluorescence that was not captured by its profile. Bottom panels: schematic of a simple model of contamination (left) and how cross-talk produces spurious correlation (right). G: Top: profile of one source and six of its transient profiles, with overlaid profile outlines from this source and an adjacent source. Bottom: Spatially-averaged activity of this source while the mouse traversed a virtual linear track. Colored arrowheads indicate transients whose profiles are shown above.

To identify whether these transients reflect activity from the source they were assigned to, the original, motion-corrected movie is examined in the spatial and temporal vicinity of each transient. We seek to produce a time-independent summary showing which pixels contributed to the transient. The “transient profile” is defined as the best fit of the transient time course to the movie frames, producing a weighted average of the frames during the transient (see Methods). For most transients, the transient profile resembles the source profile (e.g., Fig. 1B, transients 1-4), confirming that these transients were correctly attributed to the source. For other transients, however, the transient profiles bear no resemblance to the source profile (e.g., Fig. 1B, transients 5-8), indicating that they arose from a different source and were assigned in error. These two types will be respectively referred to as “true transients” and “false transients.”

To obtain a more complete picture of true and false transients, a human expert examined all 33,090 transients from 1,325 sources in this movie (for representative examples see Supp. Fig. 1–4). Most transients could be confidently classified as either true (67%) or false (13%). Other transients (20%) were not simply true or false, and fell into more complex categories; these were omitted in the initial analysis but will be considered below. Most identified sources (66%) contained at least one false transient, and for some sources (9%) the majority of transients were false (Fig. 1C, top). Additionally, significant contamination was also observed when counting spikes that were inferred from true or false transients, despite deconvolution under-emphasizing the smaller false transients (Fig. 1C, bottom). The amplitudes of true and false transients were generally comparable, both when pooling across all sources (Fig. 1D) and when comparing amplitudes within each source (Fig. 1E), though usually all of the very largest transients were classified as true.

False transients arise when a source profile overlaps with fluorescence that is not explained by any of the identified sources. This could be because some sources were not identified during cell finding, or because the spatial extent of a source was not perfectly captured by its profile, leaving a region of residual fluorescence that bleeds into other sources.

To show how false transients can affect scientific conclusions, we briefly consider two cases. First, when the activity of one source bleeds into another, it can produce a spurious correlation (Fig. 1F). In this case, the profile of one neuron (source 2) does not fully capture its true extent, and some of the uncaptured fluorescence (yellow arrowhead) overlaps the soma of another neuron (source 1). In the frames shown, only source 2 is active, but the estimated time courses show activity in both sources, creating a nonzero Pearson correlation in this time range (r = 0.73, p < 1e-35). A simple simulation (Fig. 1F bottom) demonstrates that even a small amount of bleed-through can create a significant correlation between two independent neurons.

The second case of contamination changing a scientific interpretation occurs in a CA1 place cell [27]. Place cells are typically active in a small subset of an environment, in this case a virtual linear track [10], and an important question in understanding place coding is how many discrete place fields are maintained by each cell [13]. On most traversals, this cell appears to be reliably active in two locations (Fig. 1G, blue and red arrowheads). But examining transient profiles shows that activity at one location is due to false transients (red arrowheads), and thus an observer would incorrectly ascribe two fields to this cell rather than one.

Though these results demonstrate that false transients can occur and interfere with scientific conclusions, the prevalence and significance of contamination are likely to be specific to each application, and thus we cannot draw conclusions about its occurrence in general. Instead, these examples illustrate the need for tools that allow users to identify and correct false transients in their own data, ideally tools that are compatible with a wide range of cell finding and post-processing algorithms.

### 2.2 Quantifying contamination

We begin by developing a quantification to automatically determine whether transients arose from their assigned source or resulted from some other process. Our approach is based on an assumption that is central to essentially every algorithm for calcium imaging analysis: movies are space-time separable. That is, each movie frame should be equal to the sum of all source profiles, weighted by their respective time course values. A corollary of this assumption is that the profile of each source should resemble the bright pixels during a true transient, but not during a false transient. (The more complex case of overlapping true and false activity will be considered later.)

Thus we assess each transient based on the shape of pixels that are active, which can often be identified by examining the movie itself (Figure 2B). Manual inspection of each frame, however, can be prohibitively time-consuming, and is difficult for small transients close to the noise level. Therefore we seek to develop a spatial summary that preserves as much information as possible about which pixels contributed to the transient, while also maximizing the signal-to-background ratio when collapsing over time.

**Figure 2:**
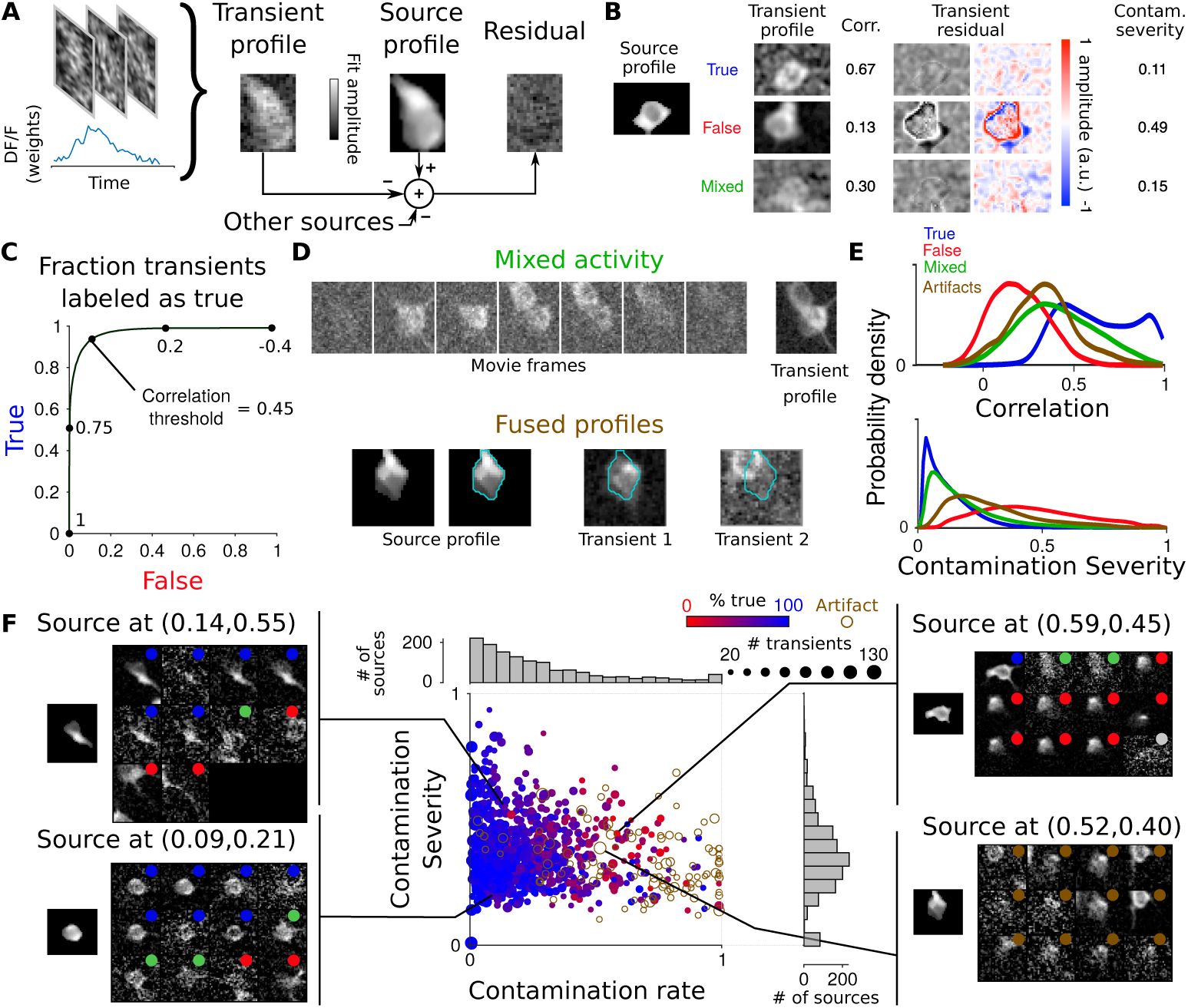
Automated evaluation of transients. A: Schematic of how the transient profile and transient profile residual are computed. B: Three transients (rows) for one source, source profile shown at left. Shown for each transient are the human classification (true, false, or mixed), the transient profile with Gaussian blurring to highlight structure (radius 1 pixel), the correlation metric value, the transient residual (shown with two colormaps), and the contamination severity metric. C: Performance of automatic classification using the correlation metric. D: Top: example of a transient with mixed activity (movie frames shown as three-frame averages). Bottom: example of a fused profile. Left images show source profile with and without contour, and right images show two representative transient profiles with contour overlaid. E: Distribution of correlation values (top) and contamination severity (bottom) for sets of transients defined by human classification. F: Center: FaLCon plot for 1,325 sources identified in one CA1 movie (same dataset as shown in Fig. 1A) at a correlation threshold of 0.5. For axis definitions, see text and Methods. Each point is one source, color shows human classification. Left and right columns: For each of four example sources located at different regions of the FaLCon plot, images show source profile (single panel) and all transient profiles (grouped panels). Inset dot colors show human classification of each transient (gray indicates transients that could not be confidently classified).

First, consider the simple average of frames during a transient. Though the noise amplitude is reduced compared to single frames, this procedure loses temporal specificity by treating all frames equally. We instead exploit the time course information and weight each frame by the transient amplitude. The weighted average provides better denoising and reduces the presence of unrelated transients from nearby sources (Figure 2B).

Using the weighted-average definition of transient profiles, we seek to automatically classify each transient as true or false. Consider the following model:

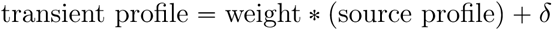

In a true transient, the weight is positive and *δ* is random noise. In a false transient, the weight is zero and *δ* is noise plus a different source. A simple way to distinguish these cases is to test whether the weight is large relative to δ, for example by testing whether pixels of the transient profile and source profile are linearly related. This test reduces to an intuitive procedure: checking whether the Pearson correlation between the transient and source profiles exceeds a certain threshold. This correlation-based metric provides good separation of true and false transients (Figure 2B, 2C), where ground truth is given by human expert classification.

The plotted performance, however, only includes transients judged by a human expert to be clearly true or false. More complex transient types can also occur, such as mixed transients, which contain a combination of true and false activity (Figure 2D, top). It is also possible that some estimated source profiles reflect an artifact, such as when partial shapes of two separate sources are merged into one and the “fused profile” is subsequently assigned transients from both (Figure 2D, bottom). Alternatively, a source shape can be generated by fluorescence changes arising from uncorrected tissue motion. Such possibilities will be collectively categorized as “artifact sources”.

How should automatic classification be designed to handle these more complex cases? As transients from artifact sources reflect an erroneous record of activity, they should be classified as false. For mixed transients, however, the answer is not as clear. If an observed transient is the sum of true and false activity, it is impossible to capture just the true component with any accept-or-reject classification scheme. With this ambiguity in mind, we will nevertheless err on the side of caution and aim to classify mixed transients as false, since their activity is at least suspect, if not an outright error.

Fortunately, the correlation metric values of mixed transients and artifact sources are largely similar to those of false transients (Figure 2E). Their distributions include a thin tail of high correlations, however, indicating that sometimes mixed and artifact transient profiles strongly resemble the source profile. This observation might suggest development of an alternative metric, such as testing a null model that pixels of *δ* are spatially independent. One implementation of this idea, based on a two-dimensional extension of the Ljung-Box quantile (sLBQ) test, performs better at rejecting artifacts, but the benefit is outweighed by lower overall performance (Supplementary Figure 6).

Instead of devising a new metric, we will overcome these limitations by supplementing the correlation metric with a secondary quantification. First, we define for each source the *contamination rate*, equal to the fraction of transients classified as false by the correlation metric. We then define the *contamination severity*, which quantifies how far false transient profiles deviate from the source profile. It is calculated as the ratio of unexplained fluorescence to total fluorescence: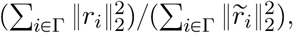 where *r*_*i*_ is the *i*^*th*^ transient profile residual, 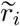 is the *i*^*th*^ transient profile with other source profiles removed, and Γ is the set of transients marked as false (see Methods). The severity allows the various causes of false classification to be distinguished (Figure 2B, E).

One application of contamination rate and severity is to decide whether scientific effects might have been caused by contamination. For example, the degree to which an effect is present in each source might be compared to these measures of contamination.

But our primary application is to provide an efficient visual summary of many sources. Consider a plot comparing the contamination rate and average contamination severity for each source (Fig. 2F). Each region of this plot supports a particular conclusion about the quality of source estimation. The lower-left quadrant contains sources whose shape and time course are largely accurate. The lower-right quadrant indicates a subtle mismatch in estimating the source profile, implying the time course is generally free of contamination. Any sources located in the upper sections of the plot, however, likely suffer from contamination, and the horizontal axis indicates whether contamination is rare (left) or frequent (right). These metrics are in approximate agreement with human expert classification, indicated by the color of each point. Contamination can also be estimated using the sLBQ test (Supplementary Figure 7). In this case, the utility of contamination severity to distinguish shape mismatches is more readily apparent, but true and false transients are not as well separated overall.

In addition to showing the variability of sources within a dataset, this plot can also be used to compare across datasets. Consider equivalent plots depicting the sources obtained from two other movies, one of CA1 tissue densely labeled with a different calcium indicator (Supplementary Figure 8), and one of sparsely labeled cortical neurons (Supplementary Figure 9). Compared to the dataset shown in Figure 2F, it is immediately clear that the second CA1 dataset suffers from even more extreme contamination, while the cortical dataset has little to no contamination, impressions that are also validated by human classification. This visualization, which we refer to as a Frequency and Level of Contamination (FaLCon) plot, thus provides a bird’s eye view of the degree of contamination in an entire dataset.

### 2.3 Geometric basis for contamination filtering

While we do not attempt to make a general statement about the prevalence or consequences of false transients beyond the data analyzed here, its occurrence demonstrates the need for a method to eliminate or reduce contamination. One simple approach might entail manual classification to reject false transients, and we have developed software tools to expedite this process. Nevertheless, this procedure suffers from several limitations. First, human classification is not reproducible, and in the case of large datasets it can be prohibitively time-consuming. Second, transient-based classification can only be performed when activity is sufficiently sparse that individual transients can be distinguished and analyzed separately. Finally, a keep-or-reject classification scheme cannot resolve cases when true and false activity occur at the same time, a circumstance of particular importance when considering joint statistics of neural activity.

To address these limitations, we develop an automated method for filtering out false activity that does not rely on parsing time courses into discrete transients. Our approach is based on explicitly modeling contamination so that it can be removed during time course estimation at the resolution of individual frames and pixels. Before introducing formal definitions, we first provide a geometric intuition for how false transients arise, as well as the motivation for our proposed solution.

Consider a small, two-pixel movie (Figure 3A). A frame in this movie can be represented as a single point in 2D space, and a source profile can be represented as a line passing through origin. (The profile is a line, rather than a point, because scaling its amplitude does not change its shape.) A frame in which the source is active (i.e. a true transient) is a point that lies close to the line (B), while a frame from a different source (i.e. a false transient) is a point far from the line (C).

**Figure 3:**
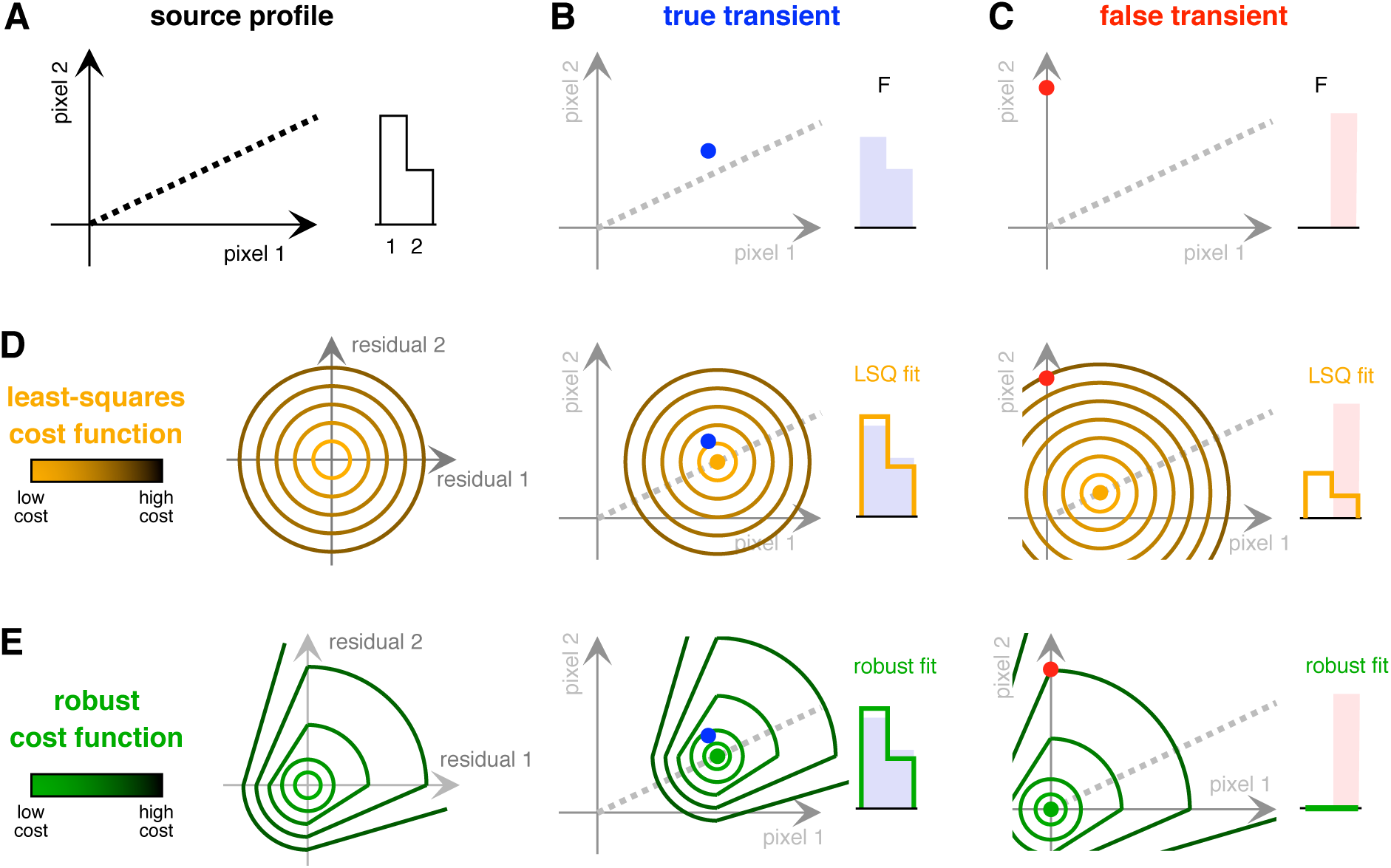
Geometric intuition for proposed solution. A: Visualization of a simulated source profile in a two-pixel movie. Left: dashed line shows points in 2D space corresponding to all possible scalings of this profile. Right: Brightness vs pixel number for one scaling of this profile. B: True transient represented as a point (left) and a plot of brightness (right). C: Equivalent plots for a false transient. D: Left: Contour lines of the least squares cost function. Middle and right: best fit of the source profile (gold point and gold outline) using least squares. E: Left: Contour lines of a hypothetical robust cost function. Middle and right: best fit of the source profile (green point and green outline) using the robust cost function.

Assuming the noise in each pixel is *i.i.d.* Gaussian, the optimal method to estimate activity is to find the scaling of the profile that minimizes the summed squared error. For a two-pixel residual, squared error is a parabola centered on the origin, and thus its contour lines form concentric circles (D, left). Geometrically, identifying the least-squares solution is equivalent to placing the concentric circle contours on the fit, then moving the fit back and forth until the cost of the data is minimized (D, middle and right). For a true transient, this method produces a good fit. But for a false transient, the energy of the other source pushes the fit to a nonzero estimate of activity.

One approach to achieve robustness is to be more permissive of positive residuals [15], since the movie is expected to contain fluorescence that arises from unknown sources. An example of such a cost function is depicted in Figure 3E (left). It is essentially a skewed version of the least squares residual, in which large positive residuals have a lower cost, while small residuals, or large negative residuals, have the same cost as least-squares. For true transients, this cost function produces results identical to least squares (E, middle), yet it also prevents interference from false transients (E, right).

This simple diagram illustrates the fundamental principle underlying our approach. But it does not fully specify the cost function, since there are many ways to permit large positive residuals. Moreover, focusing simply on this desired outcome would leave out useful *a priori* knowledge of the physical processes that generated the data. We therefore approach the problem of robust time course estimation by developing a generative model to describe contamination explicitly. As we will show, the desired statistical properties emerge naturally from the model, and allow us to retain an intuitive connection to the underlying physical processes.

### 2.4 SEUDO model and algorithm

Most methods for estimating time courses assume a straightforward model: observed fluorescence is equal to source profiles multiplied by source time courses, plus noise (Fig. 4A). Our model adds an extra term to account for unexplained fluorescence, which includes sources not identified during cell finding, as well as sources whose spatial extent was not captured by their estimated profiles. Unexplained fluorescence is modeled as a sum of narrow Gaussian kernels whose amplitudes are *i.i.d.* exponential. Together, these kernels can potentially sum to any shape of contaminating fluorescence.

**Figure 4:**
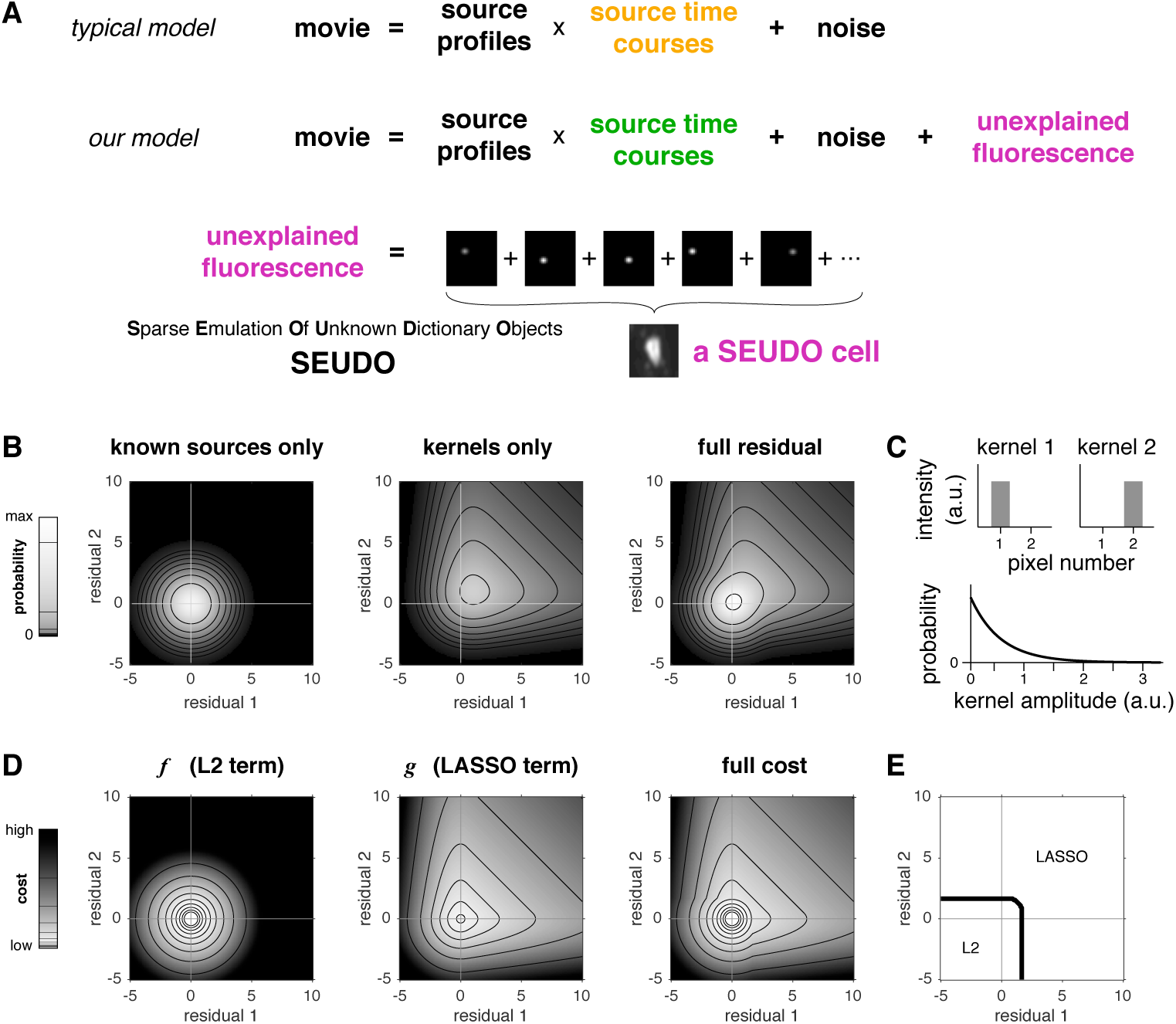
Explicit model of unexplained fluorescence. A: Schematic of two models for calcium imaging data. In a typical model, source time course weights (gold) are estimated using only known profile shapes. In our model, time course weights (green) are estimated using a procedure that accounts for unexplained fluorescence (magenta). B: Calculated probability distribution of the SEUDO residual (unexplained fluorescence plus noise) in the case of a two-pixel movie. Left images show individual components, right image shows full residual. C: Gaussian kernel spatial profiles (top) and amplitude distribution (bottom). D: Calculated cost for the objective function defined by SEUDO. Left images show individual terms, right image shows full cost. Right: boundary between the *ℓ*_2_ term and the LASSO term (*f* and *g* in Eq (2)) being lower cost.

We refer to this model as Sparse Emulation of Unknown Dictionary Objects (SEUDO). The collection of Gaussian kernels, referred to as a SEUDO cell, serves to absorb unexplained fluorescence, and thus prevents contamination from interfering with identified sources. In the remainder of this section, we develop the model formally and use it to derive a robust estimator of activity for each source.

Mathematically, we denote the observed fluorescence image in frame *t* as *y*p*t*q, such that

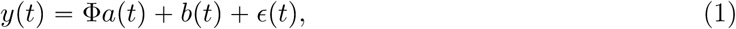

where *a*(*t*) is the vector of activation levels for known sources, Φ denotes their spatial profiles, *b* (*t*) is the image of structured contamination from other sources, and *c*(*t*) is the unstructured observation noise vector. The model considers each frame *t* independently, and so time references will be omitted. We formulate the inverse problem of finding source activity *a* from the observed fluorescence *y* as a maximum likelihood estimation, i.e. maximizing P *y*|*a* with respect to *a*. This formulation enables us to explicitly include activity from unknown (interfering) sources *b* by modeling how the interfering activity influences the observed fluorescence via a joint likelihood P_*y*_p*y|a, b*q, 2) modeling the activity of the interference via P_*b*_p*b*q, and 3) observing that the likelihood function can be obtained by marginalizing over *b*,

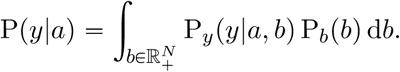

Since unexplained fluorescence is relatively rare, we consider it a special case that occurs with probability *p*. In this case, *b* is non-zero, and we model its values with the prior distribution P_*b*_(*b*). Consequently, with probability 1 *p*, there is no interference and *b* 0. Treating these cases as two components in a mixture distribution, we can rewrite the likelihood as

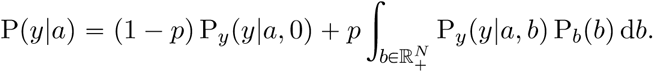

The distribution of measurement noise P_*y*_(*y|a, b*) denotes the likelihood of the observed movie frame given both the signal *a* and interference *b*. Although noise in calcium imaging movies is typically determined by the properties of photon detectors [35], and thus the variance scales with the mean, for simplicity we will approximate the noise as a Gaussian distribution with constant variance 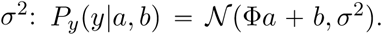. The problem is then reduced to specifying a distribution for *b* that captures the appearance and amplitude of interfering activity while also interacting elegantly with the likelihood functions to produce an efficient maximum likelihood procedure.

We posit that the interfering activity *b* should be nonnegative, spatially correlated, and spatially sparse. To attain these properties, we model *b W c* where *c* is nonnegative and sparse (i.e. mostly consists of zeros) and *W* represents a dictionary of positive 2D Gaussian kernels that induce spatial correlations. Thus *W _c_* is equivalent to the convolution of a fixed Gaussian kernel with a sparse coefficient set, essentially constructing each interfering source with a small number of kernels. Mathematically, this prior can be expressed via a single-parameter description of the Gaussian kernel (the radius) and a single-parameter exponential distribution over *c* (*i.i.d.* exponentially distributed with variance parameter λ). While more complex models might include temporal structure or higher-order spatial correlations, this model captures the basic characteristics of interfering processes and results in a much simpler estimator.

Using this model of *b*, we can derive (for full derivation see Methods) a maximum log-likelihood estimator of the form

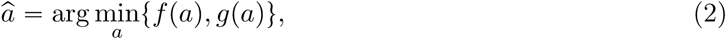

where *f* (*a*) is the maximum likelihood function conditioned on *b* = 0, i.e. the least-squares estimate 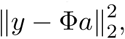 and *g*(*a*) is the value at the minimum of the LASSO estimator

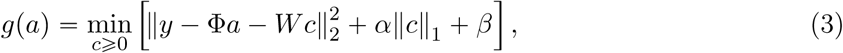

for constants *α* = 2σ^2^λ and *β* =—2 *σ*^2^ (log(1 /*p*— 1) + *N* log(1/*λ*)) where *N* is the number of pixels. The final estimate is achieved by minimizing *f*(*a*) and *g*(*a*) separately, then taking the value of *a* produced by the smaller of the two function evaluations. This estimation procedure can be thought of as two-hypothesis likelihood test to discern whether there was, or was not, unexplained fluorescence, then choosing the most likely case.

Estimating source time courses by solving this optimization constitutes the SEUDO algorithm. A significant advantage of SEUDO is that it fits the shape of unexplained fluorescence using LASSO, a well-characterized procedure for model selection, and therefore the fit can be computed efficiently using a wide array of available methods [34].

To compare SEUDO to the idealized robust cost function described above, we consider it applied to a two-pixel movie, as in Figure 3. The forward model of the residual (observed fluorescence minus the activity of known sources) is depicted separately for two cases: a frame when only the known sources are active (Figure 4B, left), and a frame that also contains unexplained fluorescence (Figure 4B, middle). When summed, these yield the full expected residual (Figure 4B, right).

Computing the maximum likelihood amounts to inverting these functions, an operation for which SEUDO is an approximate solution. To apply SEUDO in this two-dimensional case, a set of two kernels was generated (Figure 4C). Evaluating the SEUDO cost function, the L2 term *f* is the least squares cost function (Figure 4D, left), and the LASSO term *g* is skewed to be more permissive of positive residuals (Figure 4D, middle). The full cost function (Figure 4D, right) is a piecewise combination, with the boundary between *f* and *g* consisting of a simple 1D manifold (Figure 4E). Thus the cost is identical to least squares for small residuals or large negative residuals, but lower than least squares for large positive residuals. The SEUDO cost function bears a striking similarity to the idealized robust cost function described above (Figure 3E, left), consistent with rejection of false transients, and, crucially, SEUDO is derived from a generative model in which assumptions and parameters are directly related to physical processes underlying the data.

### 2.5 Parameter selection

A key factor governing the performance of SEUDO is the choice of model parameters, as these determine how much fluorescence is respectively attributed to known sources, SEUDO cells, and noise. For example, if parameters are chosen to make the cost of SEUDO cells sufficiently high, they are never used, and the estimator yields the same result as the simple least-squares solution (Fig. 5A, right). If the cost is sufficiently low, SEUDO cells are always used, and transients either suffer significant degradation or completely disappear (Fig. 5A, left). In an ideal intermediate case, false transients are absorbed by SEUDO cells, while true transients remain unaffected (Fig. 5A, middle). Our goal is to find this optimal regime.

**Figure 5:**
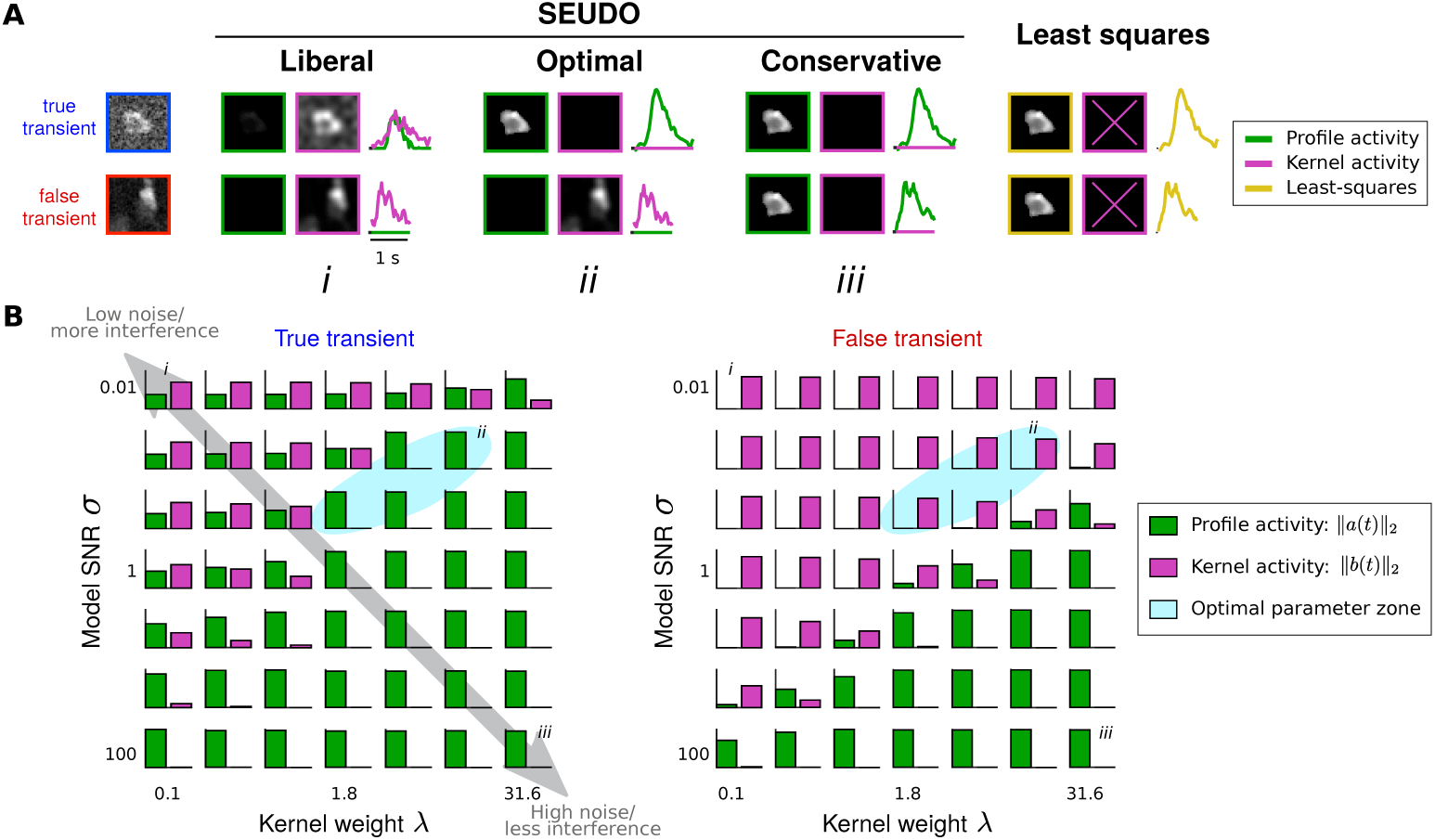
Effect of SEUDO parameters. A: Activity estimated by SEUDO and least squares for one true transient and one false transient (profiles shown at left) using several types of estimation, one type per column. Images and traces show estimated amplitude of the source profile (green) and sum of fitted Gaussian kernels (magenta). B: Sum of activity ascribed to the source profile (green) and Gaussian kernels (magenta) for the true transient (left) and the false transient (right). Each subplot shows results for one set of parameters. Roman numerals indicate parameter regimes shown in A. For blue region, see text.

The model of unexplained fluorescence contains four parameters: the radius of the Gaussian kernels, the probability of contamination *p*, the amplitude of the Gaussian kernel λ, and the noise variance σ^2^. The kernel radius determines the spatial scale of contamination, and can thus be estimated directly from examining movie frames, or chosen based on the optical point spread function and minimum neurite diameter, so its selection is not considered further.

For the three remaining parameters, inspecting the LASSO optimization in the SEUDO algorithm (Eqn. 3) reveals that they are non-identifiable, since they are combined into only two constants: *α* and β. *α* can be interpreted as a noise-to-interference ratio, i.e. the variance of photon and electronic noise (σ^2^) relative to the amplitude of unexplained fluorescence (λ). The other parameter *β* is a constant representing the likelihood that contamination occurred. It biases the choice between the least-squares and the LASSO solution by setting the *a priori* expectation of contamination (*p*) relative to the expected strength of contamination (λ), all amplified by how well such signals might be discernible from noise (σ^2^). In effect, *α* is the per-kernel cost, and *β* is the overall cost that applies whenever kernels are used.

Degeneracy allows us to explore the behavior of SEUDO by varying just two parameters. Because the estimator is relatively robust to the value of *p* (only changing when *p* flips between being close to one or close to zero), we explore parameter space by sweeping *λ* and σ^2^ (Fig. 5B). Initially we examine SEUDO applied to one true transient and one false transient from the same source (transient profiles shown in Fig. 5A).

Comparing the relative weights given to SEUDO cells (Fig. 5B, magenta bars) and the source profile (green bars) for each parameter pair, we see there are roughly two regimes. In one, SEUDO cells are employed with marked liberality (upper left), while in the other SEUDO acts conservatively, attributing even questionable activity to known sources (Fig. 5, lower right). These effects are an intuitive result of parameter selection. When σ^2^ and *λ* are both small, the model assumes low noise and high interference (recall the mean of an exponential distribution is 1/λ), implying that even small deviations from the source profile should be attributed to contamination. When both σ^2^ and *λ* are large, the algorithm assumes high noise and minimal contamination, and thus attributes most activity to the known source, since none of the fuzzy frames are expected to be a close match anyway. In between these two regions is a band where parameters are optimal (blue shaded region), and the true transient is fully preserved while the false transient is fully rejected.

To facilitate parameter selection, we provide several tools, some supervised and some unsupervised, that help to identify the optimal parameter regime for a given dataset. First, we provide a software tool that allows for rapid manual classification of transients, building up an example set that can be used to automatically identify optimal parameters, either through an exhaustive parameter search or using BayesOpt [22]. BayesOpt is an efficient supervised parameter selection tool that identifies parameters closest to the optimal set. Its central assumption is that error changes smoothly as parameters are varied, an assumption justified by the smoothly changing attribution of activity in the parameter sweep (Fig 5). The optimal parameters might vary for different sources, especially in cases where the statistics of noise and contamination are not homogenous over an entire movie frame. In this case, parameters can be specified on a source-by-source basis, or by analyzing different regions separately. Together, these tools and methods enable users to rapidly select and evaluate parameters for SEUDO.

### 2.6 pplication to CA1 data

Having developed the SEUDO algorithm and described its basic properties, we now turn to its practical application to the CA1 movie examined above, where many sources exhibited significant contamination. A key advantage of SEUDO is that it can be efficiently parallelized, and here each source is analyzed separately. Since activity in this case is also relatively sparse, SEUDO is performed only on frames in which the least squares time course exhibits a significant transient for that source. We first consider the SEUDO results using one parameter set applied to a single source (Figure 6A). In this case, SEUDO eliminated false transients nearly completely, while leaving true activity almost entirely intact.

**Figure 6:**
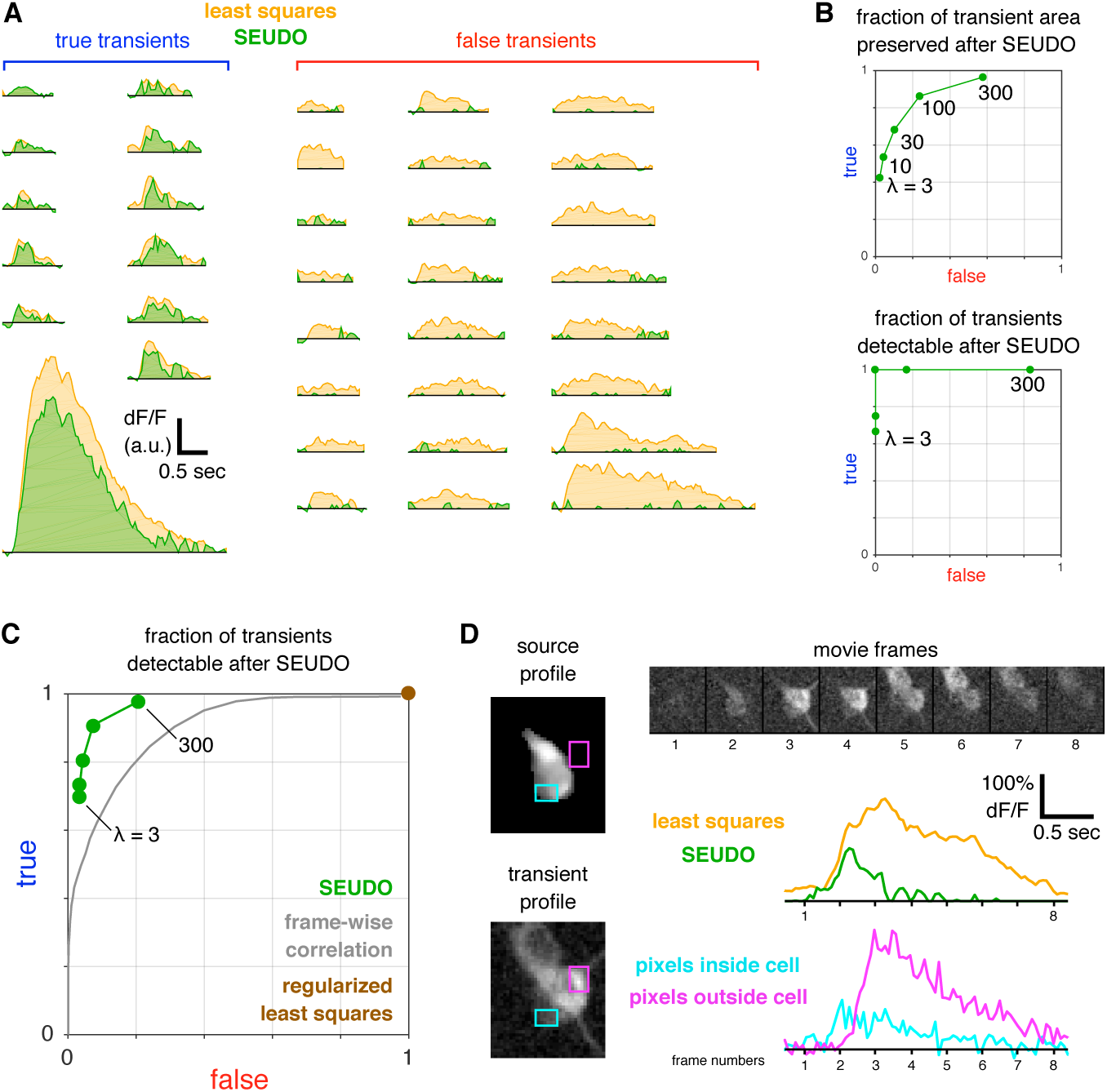
SEUDO performance on a full population of sources. A: Comparison of 36 transient time courses estimated by least squares (gold) or SEUDO (green, *λ* = 30, σ^2^ = 0.002). Within each category (human-classified true or false), transients are sorted by duration. B: Summary of SEUDO performance for five lambda values when applied to the transients shown in A. C: Performance of SEUDO (green), a simple frame-wise correlation metric (gray), and regularized least squares time course estimation (brown). SEUDO was performed using the same parameter sets as in B. The correlation metric was applied using 101 correlation thresholds ranging from -1 to 1 (see Methods). The regularized least squares method was CNMF. D: Detailed analysis of one transient with overlapping true and false activity. Shown movie frames are downsampled 10-fold to highlight when different cells become active. Cyan and magenta traces are the mean pixel value in the respective rectangular regions shown at left. SEUDO time course was computed using the same parameter set as in A.

In cases where perfect discrimination is not attained, however, there is necessarily a trade-off between removing false transients and preserving true transients. Depending on the particular needs of subsequent scientific analysis, it might be more important to achieve one of these aims over the other. To test how the SEUDO algorithm can be tailored for specific applications, we vary *λ* over two orders of magnitude while holding σ^2^ constant, and for each parameter pair compute the fractional area of preserved true and false transients (Figure 6B, top). When *λ* is small, false transients are completely removed, though at the cost of losing significant area from true transients. Conversely, large values of *λ* fully preserve true transients, though leaving about half the area of false transients.

For some scientific analyses, a more relevant metric than integrated transient area is whether or not transients are detectable, i.e. discriminable from noise. Applying the same transient detection algorithm to both SEUDO and least-squares estimation, SEUDO achieves perfect discrimination for this source, removing all false transients while retaining all true transients (Figure 6B, bottom).

To investigate the effectiveness of SEUDO more broadly, the same parameter sets were applied to the entire movie, in which 26,327 transients from 1,149 sources could be confidently classified as true or false by a human expert. Though no value of *λ* produces perfect discrimination in the full dataset, the parameters do yield similar skew towards true preservation or false rejection, showing that parameter effects generalize across the population. Using different values of σ^2^ produced nearly identical separation of true and false transients, though the range of *λ* values was different in each case (Supplementary Figure 12).

To compare these results to a baseline level of performance, we also attempt to decontaminate source time courses by applying a much simpler frame-wise algorithm, one based on a Pearson correlation criterion (see Methods). When counting detected transients, both algorithms produce a range of results that can be biased towards either true preservation or false rejection, depending on parameter choice, but SEUDO provides superior separation of true and false transients (Figure 6C). Interestingly, if performance is instead measured by the preserved transient area (Supplemental Figure 11), frame-wise correlation performs more similarly to SEUDO. These results can be reconciled by observing that transient area can be a misleading evaluation metric, given that even a significant reduction in transient amplitude can still preserve overall shape and duration (e.g. Figure 6A). We also emphasize that the correlation criterion can not distinguish the contributions of true and false transients when they occur simultaneously, a limitation that extends to any accept-or-reject classification scheme.

A unique advantage of SEUDO is the ability to model how true and false activity overlap within a single frame, allowing their respective contributions to be distinguished. This capability is shown for one example transient (Figure 6D). Examining individual frames, the source was initially active (frame 2), but before its activity completely decayed two other, overlapping cell bodies also became active (frames 3-5). The time course estimated by least squares includes both the early activity of the source as well as the late activity of contamination, yielding a transient with a mixture of the various cells, as seen in the transient profile. The SEUDO time course, however, provides a more accurate estimate of when the source was active, as verified by selecting small subsets of pixels that contain the source but not contamination (cyan), and contamination but not the source (magenta). This example illustrates that classification of single transients or single frames, no matter how it is performed, is not capable of accurately extracting source activity when it coincides with contamination. Instead, a mix of true and false activity can only be untangled by explicit modeling, such as implemented in the SEUDO algorithm.

In a final application of SEUDO, we revisit the contamination that was previously shown to affect scientific conclusions (Fig. 1F,G). For the spurious correlation found in time courses from least squares (*r* 0.333 0.09, *p* 0), SEUDO removes all but a minimal trace of residual correlation (*r* 0.018 0.1, *p* 0.0005), with the largest transients displaying no cross-talk at all (Fig. 7A). For a few frames when the two cells appear to be active simultaneously with non-identical time courses (red arrowhead), manual inspection (not shown) reveals that these are due to false transients in source 2 that SEUDO also removes. In the case of a place cell with one true spatial field and one false field (Fig. 1G), SEUDO removes the false field almost completely (Fig. 7B). And even though some transients in the true field are also removed, their absence does not substantially alter the spatial tuning.

**Figure 7:**
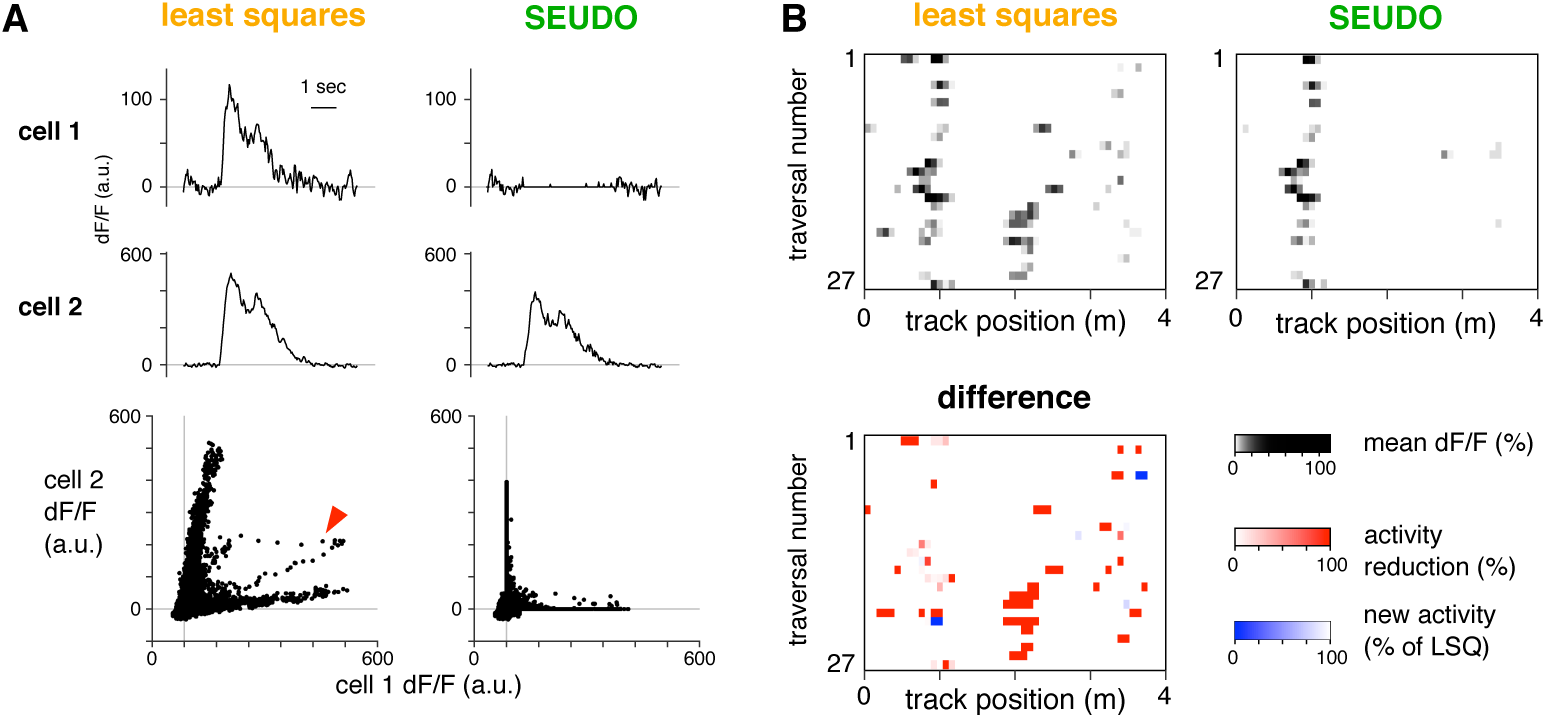
SEUDO can correct errors affecting scientific results. A: Top rows: Time courses for two simultaneously-recorded cells as estimated by least squares and SEUDO. Bottom row: scatter plot comparing time courses of the same two cells over all 41,756 frames, shown separately for each estimation technique. For arrowhead, see text. B: Top: Spatially-averaged activity for the source depicted in Figure 1G, shown separately when the time course was estimated by least squares or SEUDO. Bottom: the difference between least squares and SEUDO.

## 3 Discussion

Automated analysis methods have enabled neuroscientists to leverage the vast and rich information available in large-scale recordings. New methods, however, pose new challenges, and thus require rigorous evaluation to ensure that resulting scientific conclusions are sound. To further this aim in the analysis of two-photon calcium imaging movies, we provide a metric for identifying false transients, as well as FalCon plots to visualize the impact of contamination. Instead of relying on global statistics that can mask errors, these diagnostic tools focus on the most scientifically-relevant feature of fluorescence activity: individual transients. By aggregating results of a transient-by-transient metric into a single plot, users can appreciate at a glance the degree to which contamination affects their data.

In cases when contamination is present, we provide several avenues for addressing it. The simplest is a software interface that allows for rapid manual classification to remove false transients. While this method brings users into direct contact with data, it suffers from significant limitations: manual annotation is not reproducible, activity must be parsed into discrete transients, and overlapping true and false transients cannot be demixed. To overcome these limitations, we also develop SEUDO, a time course estimation technique to to filter out contamination. The underlying generative model in SEUDO provides a principled framework in which assumptions are made, and can be updated, explicitly, and the algorithm operates at the resolution of single frames, obviating the need to identify transients. By modeling contamination rather than classifying it, SEUDO can pull apart mixtures of true and false activity when they overlap. Importantly, SEUDO is a post-hoc analysis, meaning it can be combined with any of the growing number of cell finding procedures (e.g. [25, 11, 28, 30]). Together, these tools grant neuroscientists an informed perspective on how well the estimated time courses remain true to the underlying data, as well as the means to filter out contamination.

Some datasets are likely to be more susceptible to contamination than others, varying with factors such as labeling density, active source shapes, and optical properties. Though a comprehensive survey is beyond the scope of the methods and tools we develop here, the datasets we examine do show a trend. Greater contamination seems to occur when labeled cell bodies are densely packed and occur near major crossing dendrites (Supp Fig. 9), while contamination is less prevalent when somas are sparsely labeled and most dendrites do not travel in the image plane (Supp. Fig. 8). Our metrics will allow future studies to identify how often particular arrangements produce contamination, which could be used to inform labeling and recording strategies. It will be especially important to avoid contamination when comparing across brain regions, since apparent differences in neural coding might actually reflect differing amounts of contamination.

The problem of erroneous annotation in large datasets is not limited to imaging calcium signals. Electrophysiologists, for example, have long faced similar challenges in spike sorting, and an array of methods have been developed to quantify and correct errors [14, 36, 9, 3], many of which have become standard best practice. In two-photon imaging, signal-sources contamination has previously been described in the context of imaging dendrites [37], and one approach to removing contamination is excluding contaminated frames from further analysis, since often all spines are affected simultaneously. In large-scale movies that primarily include cell bodies, however, contamination is typically more localized, and subsequent analyses would suffer from excising whole frames, particularly when scientific questions address multicellular activity.

In the design of FaLCON plots and SEUDO, we sought to eliminate the need for excessive parameter tuning by requiring only a minimal number of parameters. For FaLCON plots, users must only specify one parameter, the correlation threshold for distinguishing true and false transients, which can be readily estimated from manually classifying a small number of sources. Similarly, SEUDO requires selecting only two parameters (aside from typical pre-processing steps, such as choosing a temporal downsampling rate). To facilitate identifying parameters optimized for the statistics of each dataset, we provide simple tools for parameter selection: ROC curve visualization, and Bayesian optimization of parameters via BayesOpt [22]. We have also sought to make execution of the SEUDO algorithm as computationally efficient as possible. Time course estimation allows complete parallelization, since the analysis is independent for each source and each frame. This also allows analysis to be restricted to a small spatial and temporal subset of the movie, reducing computational and memory overhead.

In any cell finding algorithm, estimating time courses can be performed as a separate step after source profiles have already been identified [25, 24], or as a sub-problem when optimizing for both time courses and profiles [28, 31]. In either case, essentially all methods assume a Gaussian noise model, implying they reduce to a least-squares optimization, albeit with temporal regularization terms in some cases. For this reason, we have chosen simple least-square optimization as a baseline level of performance. Though regularization does benefit time course estimation by distinguishing neural activity from noise, it cannot distinguish multiple signals when they are spatially mismatched, and thus it has a negligible impact on reducing structured contamination (Fig. 6C). Fortunately, it might be possible to combine SEUDO with other kinds of regularization because the noise model in SEUDO is orthogonal from the signal model; such priors would instantiate as modifications to *f*(*x*) and *g*(*x* in Equation (2). SEUDO could thus be incorporated flexibly into many algorithmic structures, allowing robustness to be built in to existing data-processing pipelines.

The core function of SEUDO is to distinguish whether observed fluorescence should be explained by source profiles or Gaussian kernels. This distinction might seem impossible, since kernels can construct any shape. The compressive sensing literature, however, has explored sparse estimation using over-complete dictionaries, and it provides theoretical accuracy guarantees on the LASSO estimator in such cases [23]. The guiding principal of these results is that the two structures to be de-mixed (profiles and kernels) should be oriented in sufficiently different directions in fluorescence space. Because kernels are highly localized in space, they can be accurately distinguished from source profiles, provided that the latter do not resemble single kernels. Future work might be able to derive exact guarantees and create provable bounds on SEUDO performance, increasing effectiveness of the contamination model as well as helping guide parameter selection.

In prior literature, the closest existing method to SEUDO is time-domain calcium image denoising [21]. This method provides improved temporal models of noise, and uses the one-dimensional LBQ test to assess how well the temporal auto-regressive noise model explains the data. This temporal model complements the spatial focus of SEUDO, and future work might be able to combine these ideas. Recent studies have also aimed to assess source separation by comparing spatially-localized activity within a cell profile to the full activity trace [32]. This is related to the idea of single transient assessment, but focuses on finding hidden cell activity rather than assessing the overall assignment. Again, it might be possible to combine these different modes of validation into a unified suite.

While FaLCON plots and SEUDO can assist in detecting and correcting contamination, we emphasize that our approach is only an initial attempt to address those problems, and its capabilities are not unlimited. For example, frame-wise SEUDO estimation does not include prior knowledge about the spatial and temporal properties of contamination, assumptions which might help it achieve an even closer match to human expert classification. Also, the current implementation of SEUDO does not explicitly account for activity in the neuropil [15, 16]. Through additional model development, it might be possible to overcome these limitations, as well as craft a related model to match the statistics of one-photon data. We anticipate that our framework will provide a flexible and powerful starting point for these kinds of refinements and extensions.

## 4 Author Contributions

JLG and ASC developed the analysis methods; JLG, SAK, and EHN performed the experiments; all authors wrote the paper.

## 5 Acknowledgments

We thank L. Meshulam, P. D. Rich, A. Giovannucci, and E. Pnevmatikakis for helpful comments on the manuscript.

## 6 Materials and Methods

### Experimental procedures and data acquisition

All experiments were performed in compliance with the Guide for the Care and Use of Laboratory Animals (https://www.aaalac.org/resources/theguide.cfm). Specific protocols were approved by the Princeton University Institutional Animal Care and Use Committee.

To obtain large-scale population recordings of activity, transgenic mice expressing genetically-encoded calcium indicators were surgically implanted with a chronic window that allowed optical access to the region under study. To image in CA1, the implant was designed as described previously [6], and two mouse lines were used: C57BL/6J-Tg (Thy1-GCaMP3) GP2.11Dkim/J (Jackson Labs strain 028277) expressing GCaMP3 [33], or C57BL/6J-Tg (Thy1-GCaMP6f) GP5.3Dkim/J (Jackson Labs strain 028280) expressing GCaMP6f. To image in cortex, we obtained expression of GCaMP6f by crossing several strains: Emx1-Cre (B6.129S2-Emx1^tm1(cre)Krj^/J, Jax no. 005628), CaMK2-tTA (B6.Cg-Tg(Camk2a-tTA)1Mmay/DboJ, Jax no. 007004), and TITL-GCaMP6f (Ai93; B6.Cg-Igs7^tm93.1(tetOGCaMP6f)Hze^/J, Jax no. 024103).

Activity was measured using two-photon laser scanning microscopy employing resonant galvanometer scan mirrors at a frame rate of 30 Hz. To measure spatially-modulated activity in CA1 neurons, mice were trained to run on a virtual linear track [10].

### Identifying source profiles and transient times

Motion correction and source profile identification were performed as described previously [10]. Briefly, the movie was spatially divided into a grid of 36 sub-movies and each was analyzed separately. Because of small deflections in Z, a motion artifact time course was subtracted from each source time course before filtering (median, length 10) and thresholding (zeroing points less than 4 times the robust standard deviation). Any contiguous period of above-threshold activity that lasted at least 6 frames (0.2 seconds) was considered a transient. These same steps were applied to time courses estimated by all methods considered: least squares, SEUDO, least squares decontaminated with frame-wise correlation, and regularized least squares.

### Least-squares time course estimation

Most automated calcium segmentation methods are based on estimating time courses via a regularized least-squares estimation [25, 31, 28, 30]. Mathematically, if *Y* ∈ ℝ^*N*×*T*^ is the fluorescence movie with *N* pixels and *T* frames and Φ ∈ ℝ^*N*×*M*^ is the matrix containing the spatial profiles for all *M* isolated sources, we compute the *M* × *T* matrix of per-source fluorescences *A* as

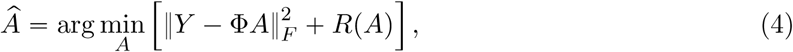

where *R A* is the regularization that imparts additional information about the expected time course structure. For example, in CNMF this regularization induces an autoregressive structure related to the calcium signal decay time-constant [31]. As contamination is essentially due to a mis-matched noise model, we seek to bypass the idiosyncrasies of each algorithm. We thus focus on time course estimation based purely on the Gaussian noise model without any regularization, i.e., least squares fitting:

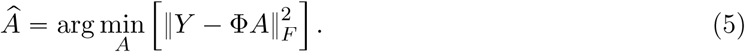

### Transient profiles

To calculate the transient profile, consider the fluorescence movie data *Y* cropped to the spatial extent of the source profile and to the temporal extent of the transient. We then seek the best representation of *Y* that has the same time course as the source’s transient activity over the duration of the event, which we denote *h*. This is accomplished by solving the optimization program

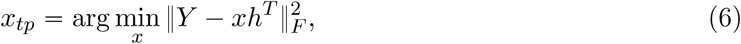

i.e., we are looking for the spatial pattern that best projects the time course into the data. The solution to this optimization program *x_tp_* is the transient profile. To see that this is simply a weighted average of the data frames, we can solve the optimization program as

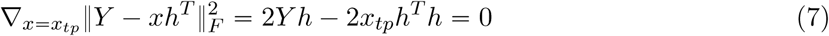

which implies that 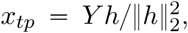, which is the weighted sum of frames where the *i^th^* frame is weighted by the *i^th^* element of the transient time course, normalized by the total magnitude of the transient event.

### Manual classification

Manual classification was performed by a human expert (author AC) based on visually comparing the shapes of source profiles and transient profiles. Transient profiles that contained the same shape as the source profile, but not shapes of other sources, were classified as true. Those containing only sources different than the one in the source profile were classified as false. Transients that looked like a mixture of true and false transients were classified as mixed. Transients in which no spatial structure was visible were left unclassified. Sources whose profile shape did not match any transient profiles (e.g. fused sources), or sources that appeared to arise from a motion artifact, were labeled as artifacts. To better resolve edge cases, small spatial blurring was at times applied to the transient profiles to highlight the shape of low-amplitude structures.

### Contamination severity

The contamination severity seeks to quantify the fraction of fluorescence not explained by the source profile. We thus define two residuals for every transient. The first is the full residual, which is the result of subtracting the contributions of all source profiles (columns of the pixels-by-sources matrix Φ) from the pixels-by-frames fluorescence movie *Y*_*ik*_ during the *i*^*th*^ transient of source *k*:

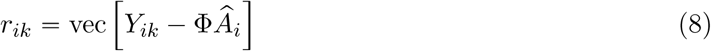

where *A_i_* is the sources-by-frames matrix of estimated temporal activity for each source and each transient frame. The second is the residual subtracting off the contributions of all the sources *except* the source profile in question, i.e., removing the column of Φ and row of *A_i_* corresponding the source the transient was assigned to:

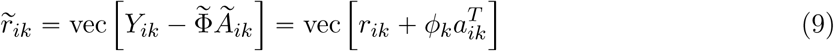

where 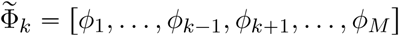 and 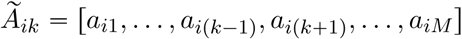 are the sets of all other sources and all other time-courses, respectively, φ_*k*_ is the profile for the *k*^*th*^ source and *a*_*ik*_ is the time course for the *k*^*th*^ source during the *i*^*th*^ transient. The contamination severity computes the amount of residual energy in all suspect transients (i.e., those that fail the correlation or LBQ test) as a fraction of the residual discounting the source

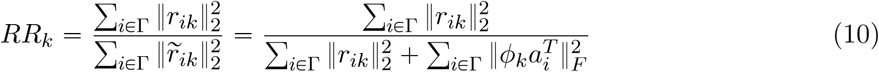

When the residual is small, and the source is active, the contamination severity is low, however when the source is not active with respect to the residual size (i.e., significant activity remains even when accounting for the source) then the contamination severity is high. To make the contamination severity more resistant to noise, simple low-pass filtering is applied (Gaussian filter with one-pixel standard deviation).

### Spatial Ljung-Box quantile test

The Ljung-Box quantile (LBQ) test is a statistical test of whether a model with *d* degrees of freedom captures the autocorrelation structure of a time series. If *x*(*t*) is the time series and *r*(*t*) is the model residual. If *x*(*t*), *r*(*t*) have length *N*, and *c*(*l*) =corr *r*(*t*),*r* (*t—l*)is the autocorrelation at lag *l*, then the LBQ test calculates the quantile statistic

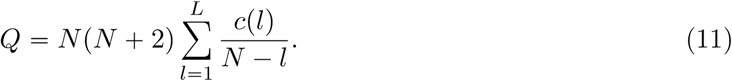

If *r*(*t*) are not correlated, then *Q* should be Chi-squared distributed with *L* degrees of freedom. Consequently, a simple quantile test can be used to determine if we can significantly rule out this null hypothesis. Specifically we test if *Q* >*q*(1 —*ζ,d*) for a given significance level ζ, which if true rules out the null hypothesis. For spatial data, i.e. *x*_*i,j*_, *r*_*i,j*_ of size *M* × *N*, we can expand the test statistic to be

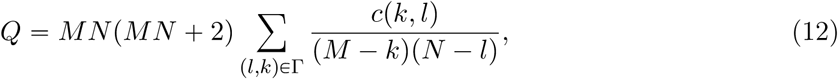

where *c_k,l_* is the spatial autocorrelation function at lags *k* and *l*, and Γ ⊆ [1, *M*] x [1*, N*] are the set of lags to be tested. With this change, the quantile test above remains, aside for adjusting the degrees of freedom to account for the increased dimensionality. For all plots in this work we set *d* to be the number of active pixels in the profile and test the set of lags between 3 and 7 lags, inclusive. This latter choice ensures that blurring from the motion correction process (which can blur neighboring pixels together) will not bias the results. Finally, we run all the tests at the *ζ* 0.01 significance level.

### Derivation of SEUDO algorithm

We seek to use the observed movie frame *y* and source profiles Φ to identify the source activations *a* (i.e. the time course value in this frame). To perform robust estimation, we assume this frame might contain structured fluorescence not explained by the known source profiles, denoted as nuisance variable *b*, which we wish to marginalize over in order to perform unbiased estimation of *a*,

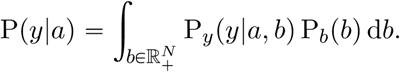

We further assume that unexplained fluorescence occurs *sparsely*, such that *b* will be nonzero with probability *p*, giving

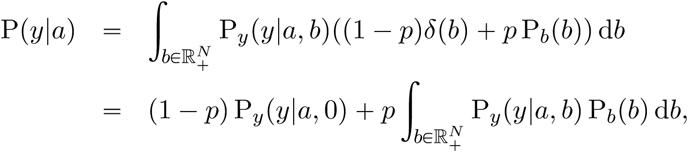

where P_*b*_(*b*) is the distribution over the set of all possible unexplained fluorescence images, and P_*y*_ *y*|a,*b*) the measurement noise distribution. Taking the noise to be additive Gaussian with mean 0 and variance σ^2^, we have

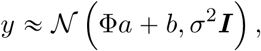

and can express the likelihood as

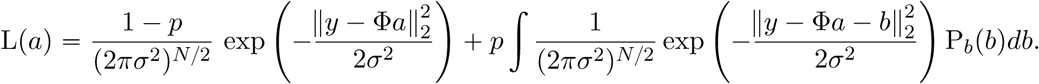

To optimize this likelihood with respect to *a*, we must specify an explicit model for the unexplained fluorescence *b*. We wish to ensure that *b* is sparse, nonnegative, and locally correlated across pixels. To achieve local correlation, we model *b* as *b*= *W_c_*, where *W* is a dictionary of 2D Gaussian kernels and *c* is a set of weights, one for each kernel. To achieve sparseness and nonnegativity, we make *c* multivariate exponential with parameter λ. For simplicity, we will also make *W* square, so that the dimensionality of *b* and *c* are the same (i.e. one kernel per movie pixel).

With the given prior over *b*, and defining the residual *r* = *y* = Φ*a*, the second likelihood term can be rewritten as

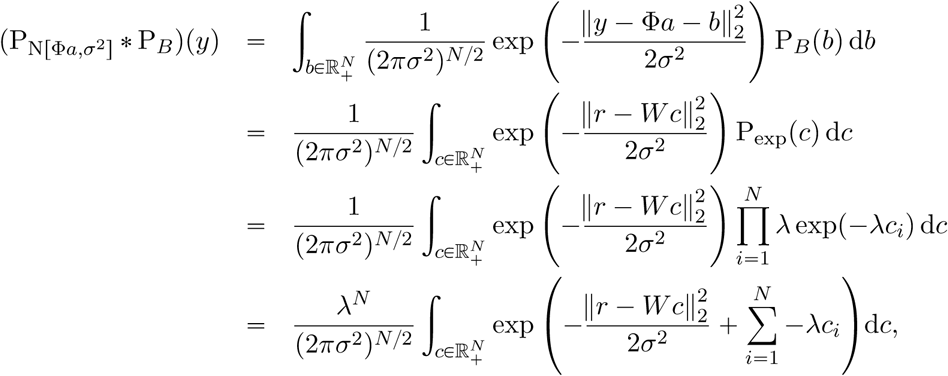

which yields the full log likelihood

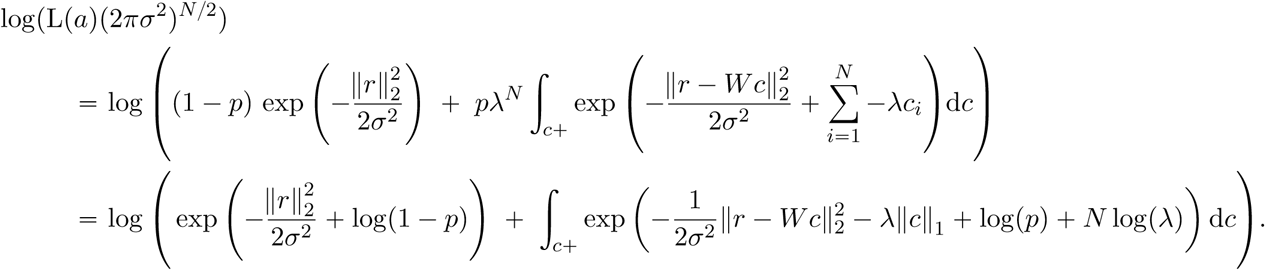

This log likelihood is essentially the log of a sum of exponentials, so we can employ the approximation

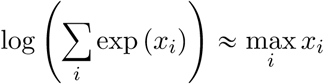

which in the limit of an integral becomes

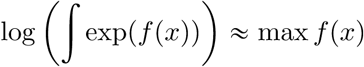

This will extremely simplify the computation of the maximum likelihood solution

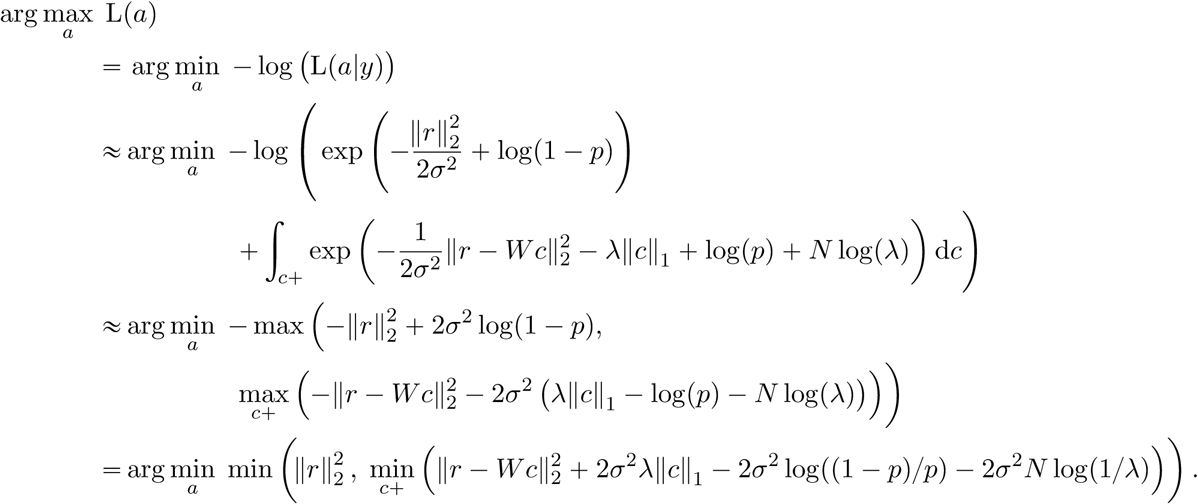

This maximum likelihood function has the form of

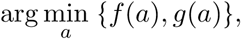

which can be solved by minimizing *f*(*a*) and *g*(*a*) separately, then using the value of *a* that yielded the smaller result. In SEUDO, *f* p*a*q_2_ is the maximum likelihood function conditioned on *b “* 0, i.e. the least-squares estimate 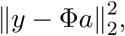, and *g*p*a*q is the LASSO estimate

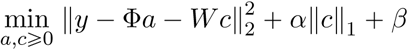

for constants α = 2σ^2^λ and *β* = —2σ^2^ (log(1/p - 1) + *N* log(1/λ)).

**Frame-wise correlation decontamination algorithm**

To generate a simple baseline for comparing to SEUDO, time courses were decontaminated using the following procedure. For each source, a small spatial subset of the movie was considered. In each frame, the Pearson correlation was computed between the movie and the source profile. If this correlation fell below threshold, the time course was set to 0. The threshold was varied from -1 to 1, and transient detection (as described above) was applied to the time course from each source.

**Transient metrics**

Two quality measures were used to evaluate the various time course estimates (least squares, SEUDO, least squares decontaminated with frame-wise correlation, and regularized least squares). No matter what time course was being evaluated, transient times were defined by the least squares time course. To summarize performance on a large number of transients, results were averaged over all true transients and all false transients separately (where true and false were defined by human expert classification).

The first measure was the fraction of transient area preserved in comparison to least squares. If 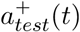 was the rectified time course estimated by the technique being tested, and 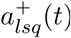 was the rectified least-squares time course, the fraction of transient area preserved was 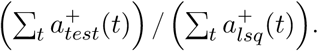.

The second measure simply evaluated whether the transient was still present, regardless of its integrated area. In this case, transient detection (as described above) was applied to the entire time course being tested. If at least one transient occurred in the period of the least squares transient, that transient was considered kept and the measure returned 1. Otherwise the transient was considered rejected, and the measure returned 0.

### Software

We created a MATLAB-based software suite that implements all the algorithms described here. The software also includes several graphical user interfaces (GUIs) to facilitate visualization and optimization of analysis parameters. For example, one GUI displays transients for manual classification (Supp. Figure 10). It allows users to classify transients based on comparing the shape of each source profile to the transient profiles, with several options for how to display them: overlaying the source profile contour, switching to high contrast colormaps, and blurring to reduce the effect of high frequency noise. Users can also interact with this GUI to specify which transients are true, false, or contain mixed activity, as well as identify artifact sources. Manual classification serves to estimate the overall level of contamination, and it establishes a training set to calibrate the parameters of automatic analysis. Other GUIs (not shown) facilitate comparing different sets of parameters to optimize performance.

All software can be downloaded from the following website: https://adamsc.mycpanel.princeton.edu/downloads.html

---

**Supplementary Figure 1:**
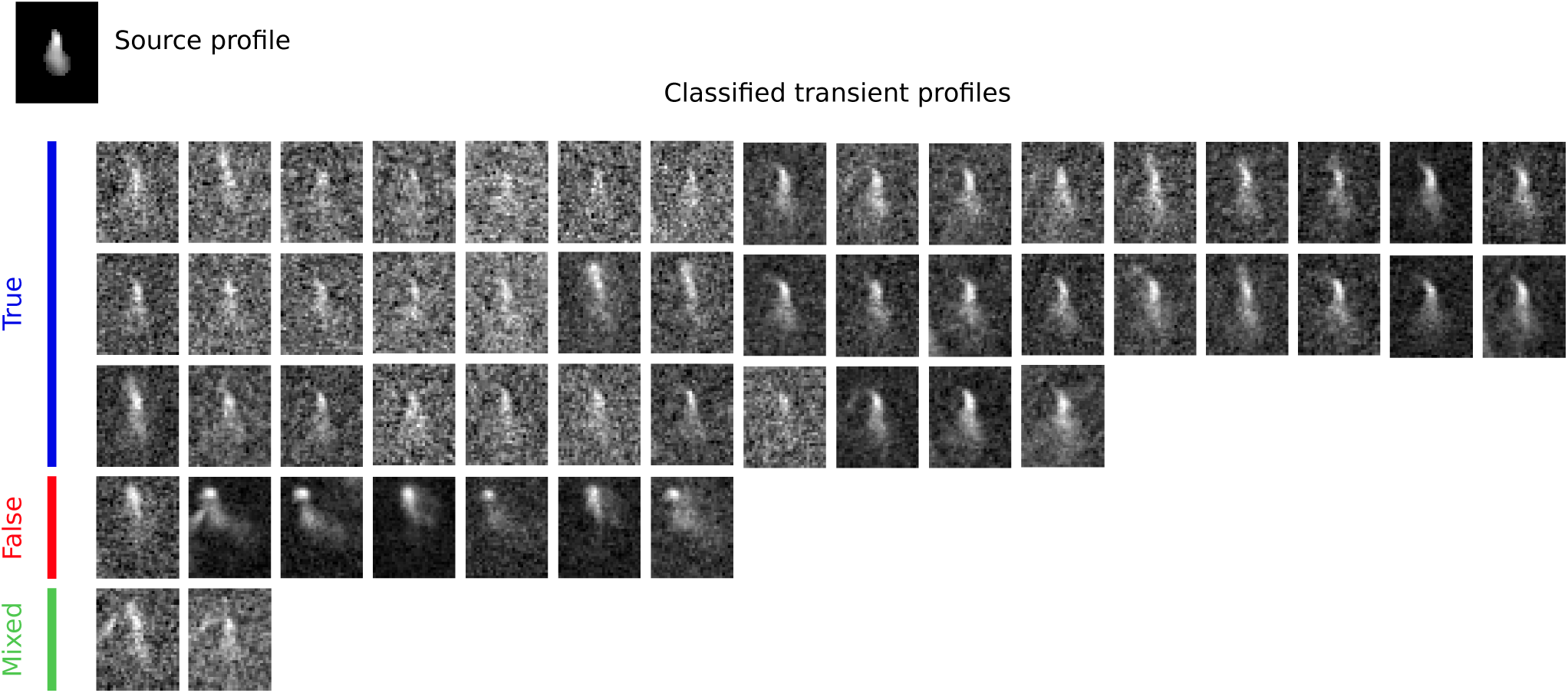
Example classified transient profiles for a single source from mouse CA1 found using CNMF.

**Supplementary Figure 2:**
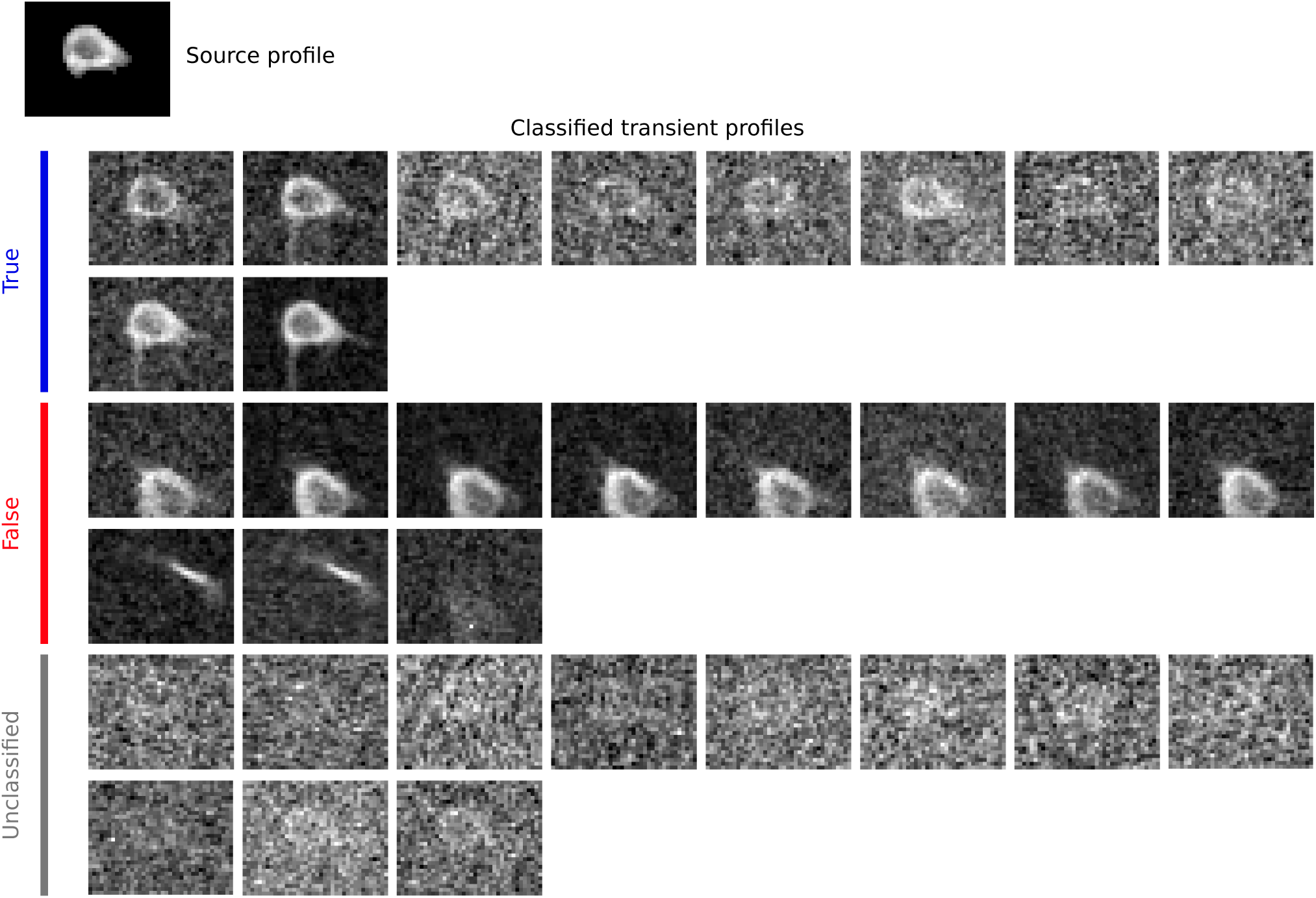
Example classified transient profiles for a single source from mouse CA1 found using CNMF.

**Supplementary Figure 3:**
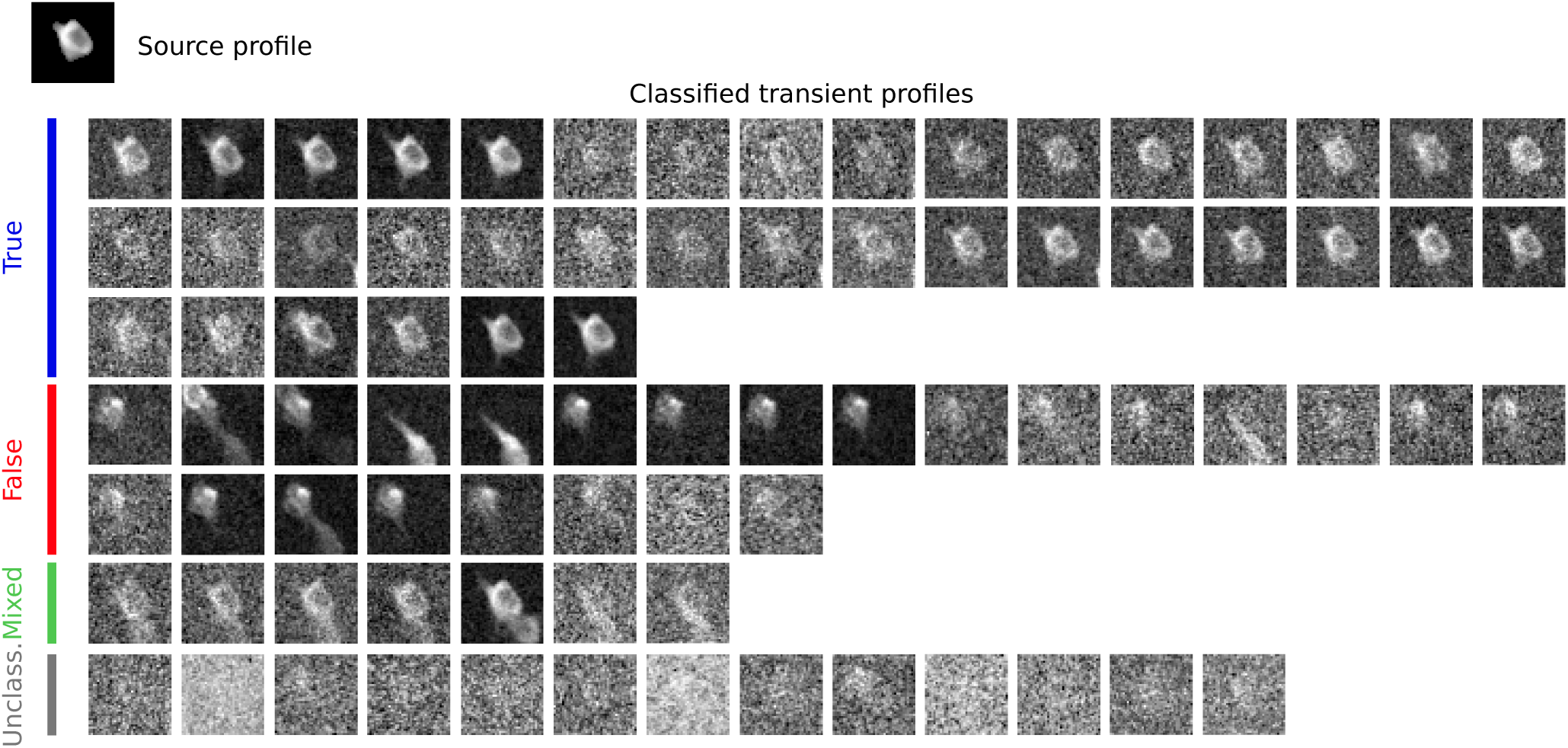
Example classified transient profiles for a single source from mouse CA1 found using CNMF.

**Supplementary Figure 4:**
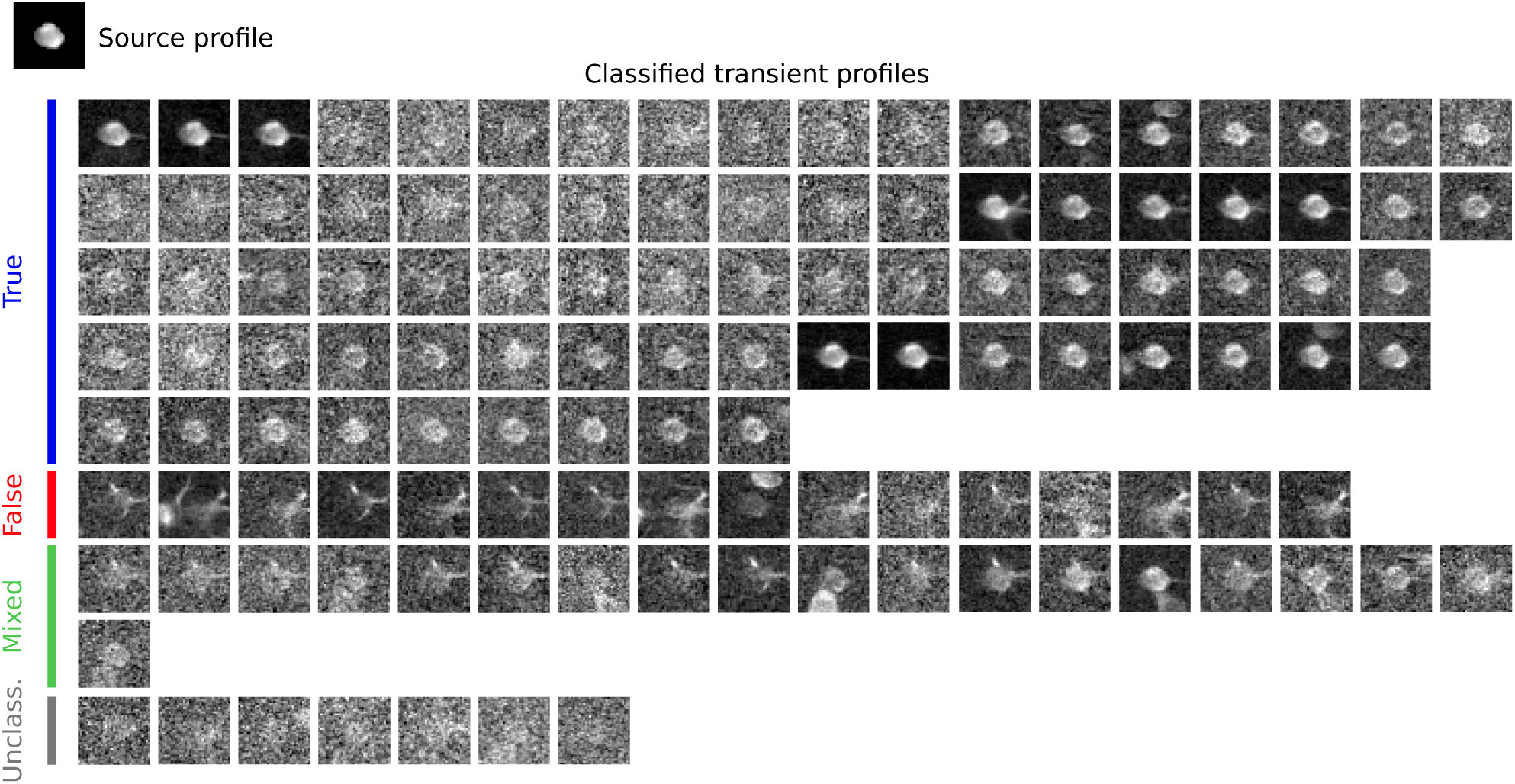
Example classified transient profiles for a single source from mouse CA1 found using CNMF.

**Supplementary Figure 5:**
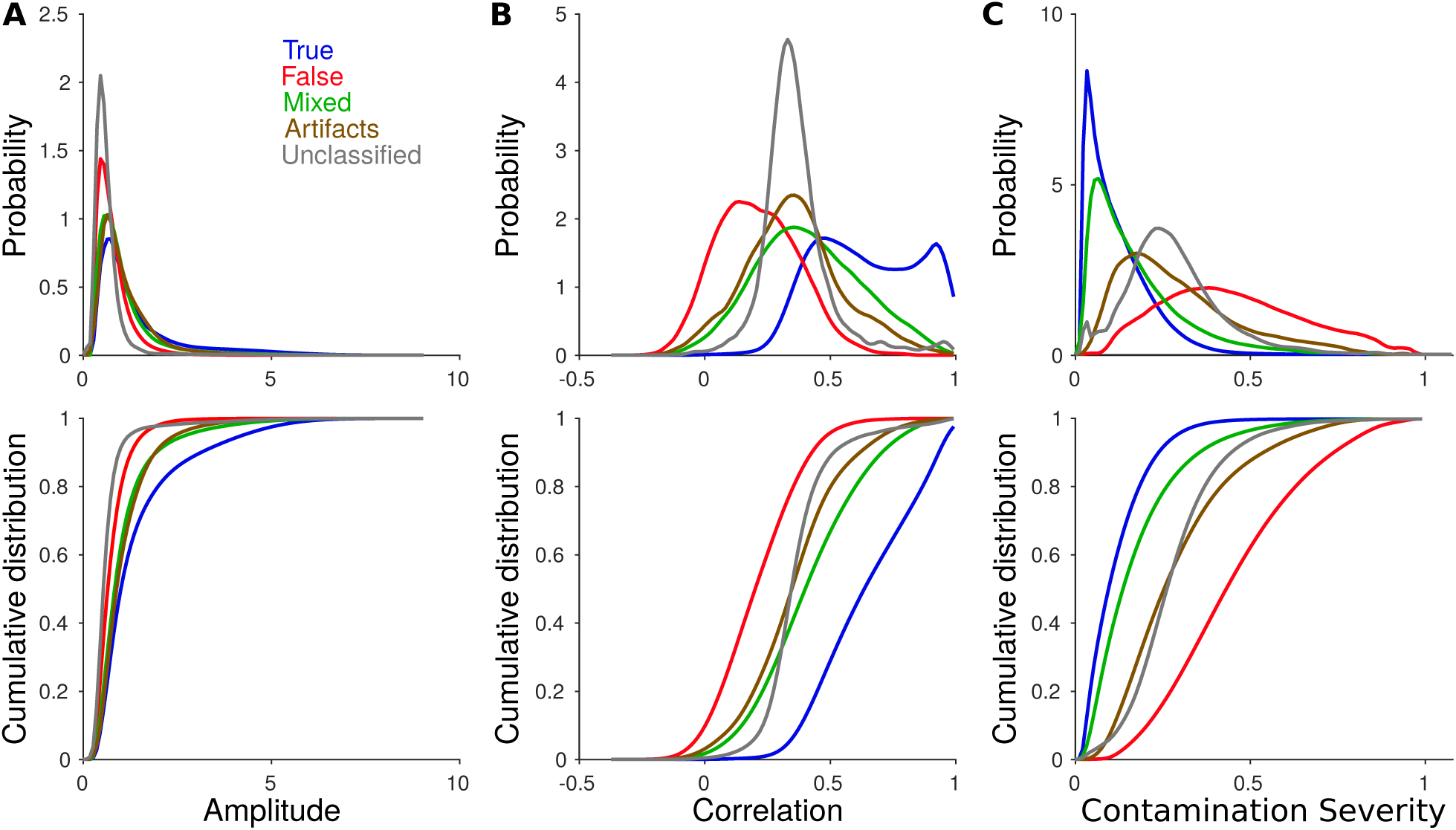
Spread of amplitude, correlation, and contamination severity for true, false, mixed, artifact, and unclassifiable transients.

**Supplementary Figure 6:**
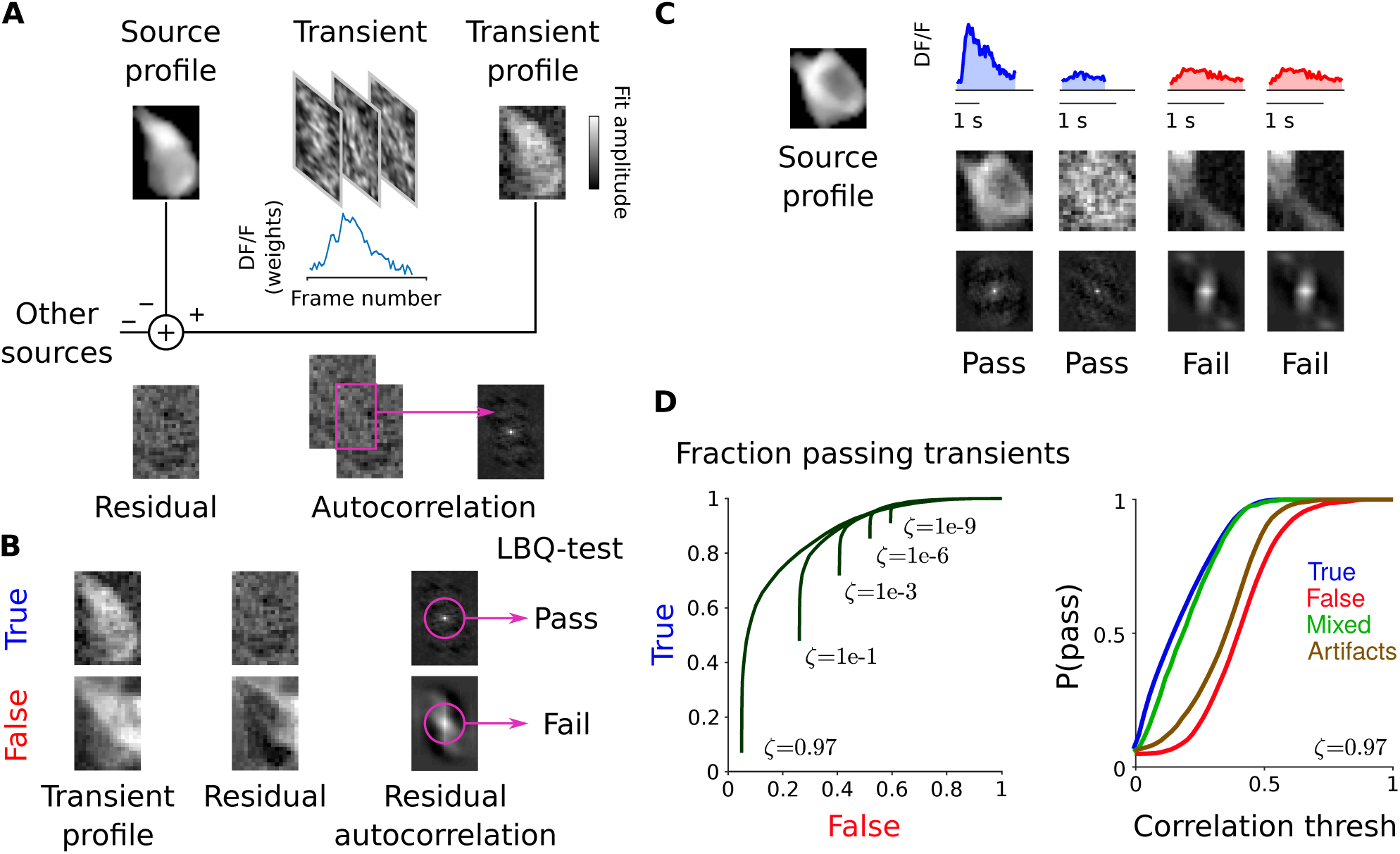
Algorithm to classify transients as true or false using the spatial Ljung-Box quartile test instead of the correlation metric. A: Schematic for how the transient profile residual autocorrelation is computed. B: Two example transients, shown to illustrate difference in residual for true and false transients. C: Classification of four example transients using the LBQ test. D: Results of the LBQ test on transients classified by human expert. Left: Results of the test applied to true and false transients for various values of α. Classification accuracy is worse than for the correlation metric (Figure 2C), likely due to due to sensitivity to other inherent nonlinearities. Right: Results of the test applied to all four transient types.

**Supplementary Figure 7:**
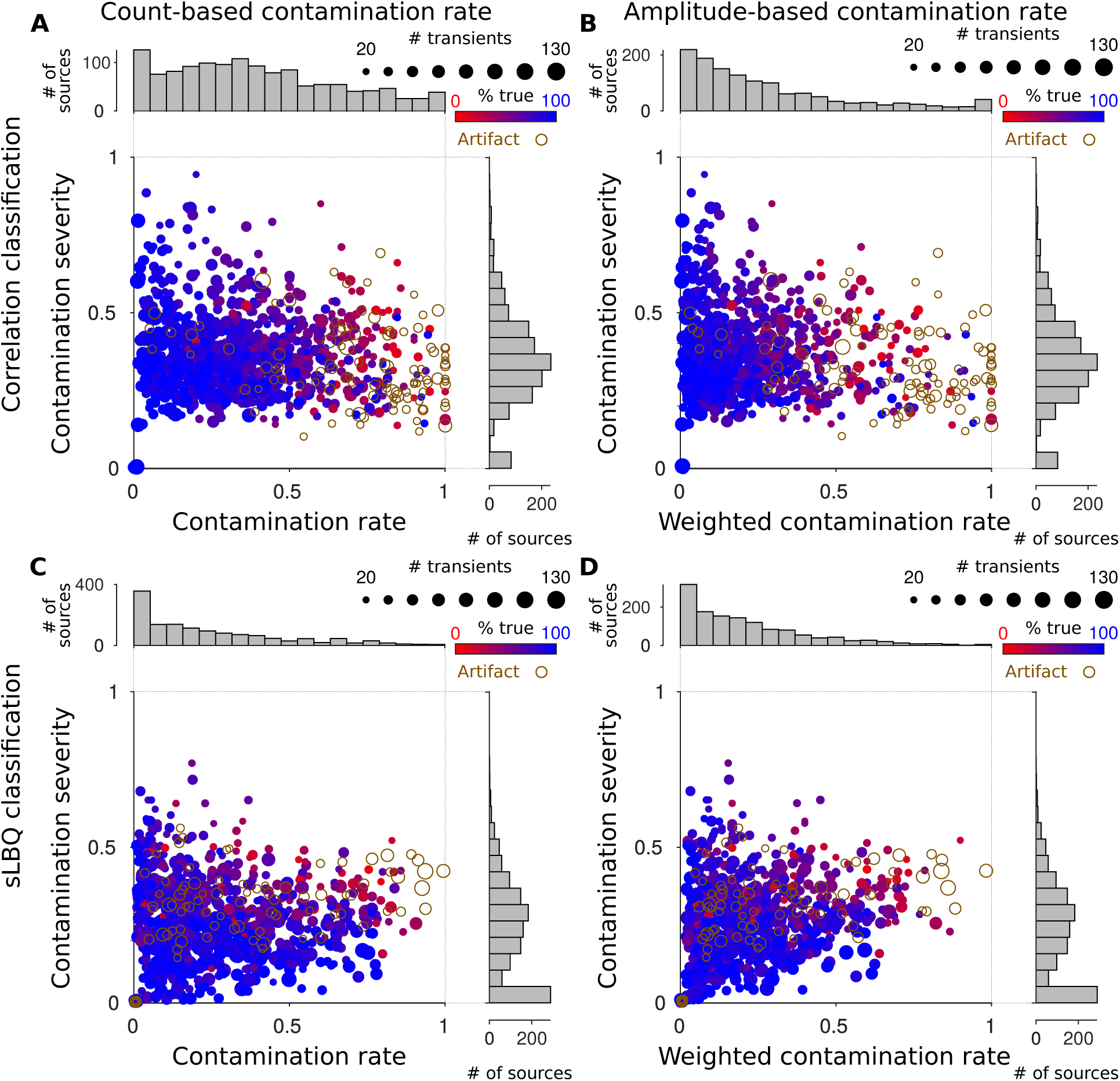
Alternate definitions of “level” and “severity” can be used to modify the FaLCon plot to have different characteristics. Two such changes are displayed here. A) A FaLCon plot using a correlation based rate estimation where the rate is the unweighted fraction of transients used. B) Same data as in A, however the level of contamination is weighted by the relative contribution based on amplitude values. C) and D) are the same as A) and B), however using the sLBQ test to determine true/false transients. As the sLBQ test is more sensitive to mis-match in spatial shapes, the level of contamination is especially important to separate data.

**Supplementary Figure 8:**
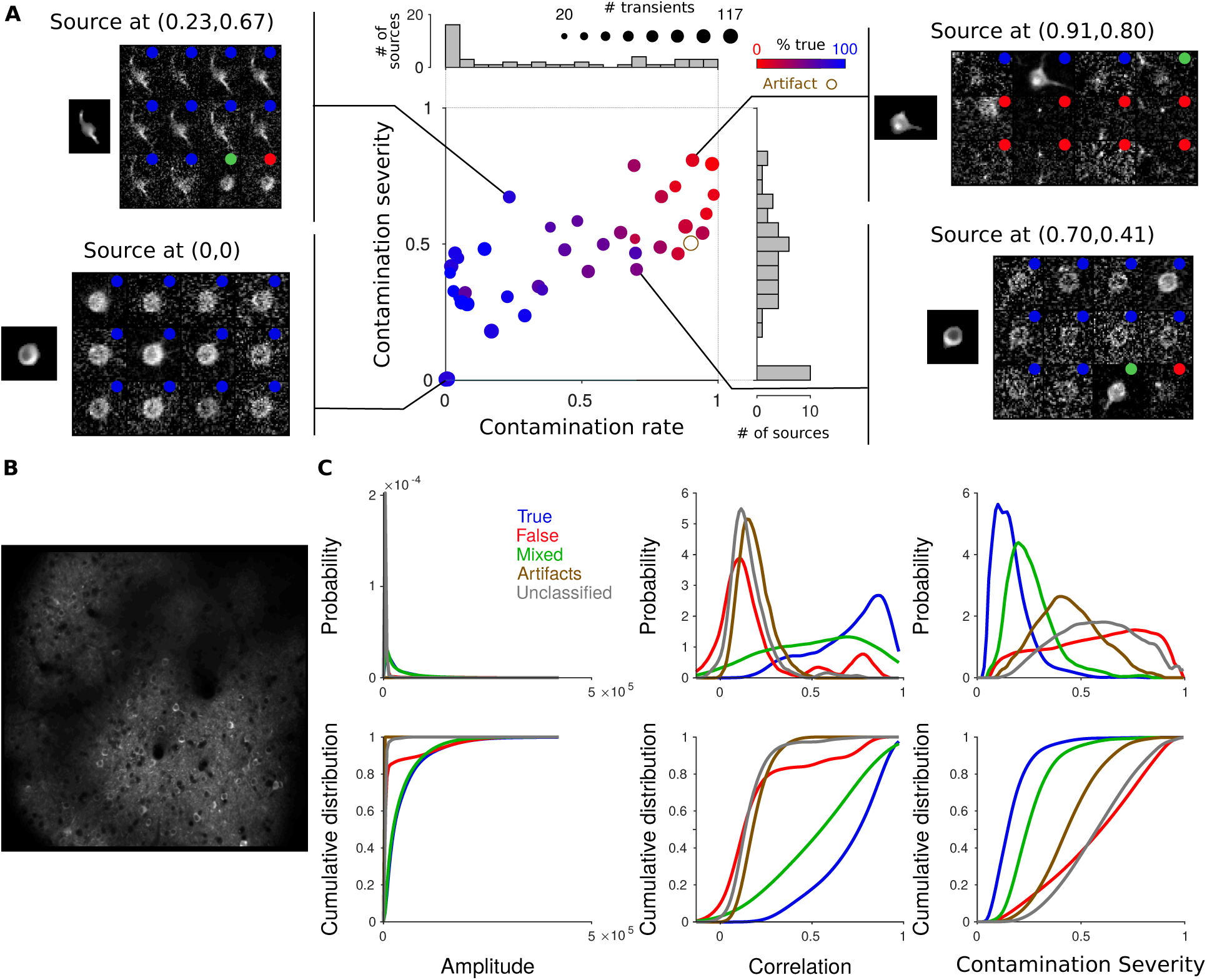
Analysis of sources estimated by CNMF from a movie of mouse visual cortical area MMA sparsely labeled with GCaMP6f. A: FaLCon plot, same conventions as in Figure 2F. B: Average movie image. C: Properties of different categories of transients, with categories defined by human expert classification.

**Supplementary Figure 9:**
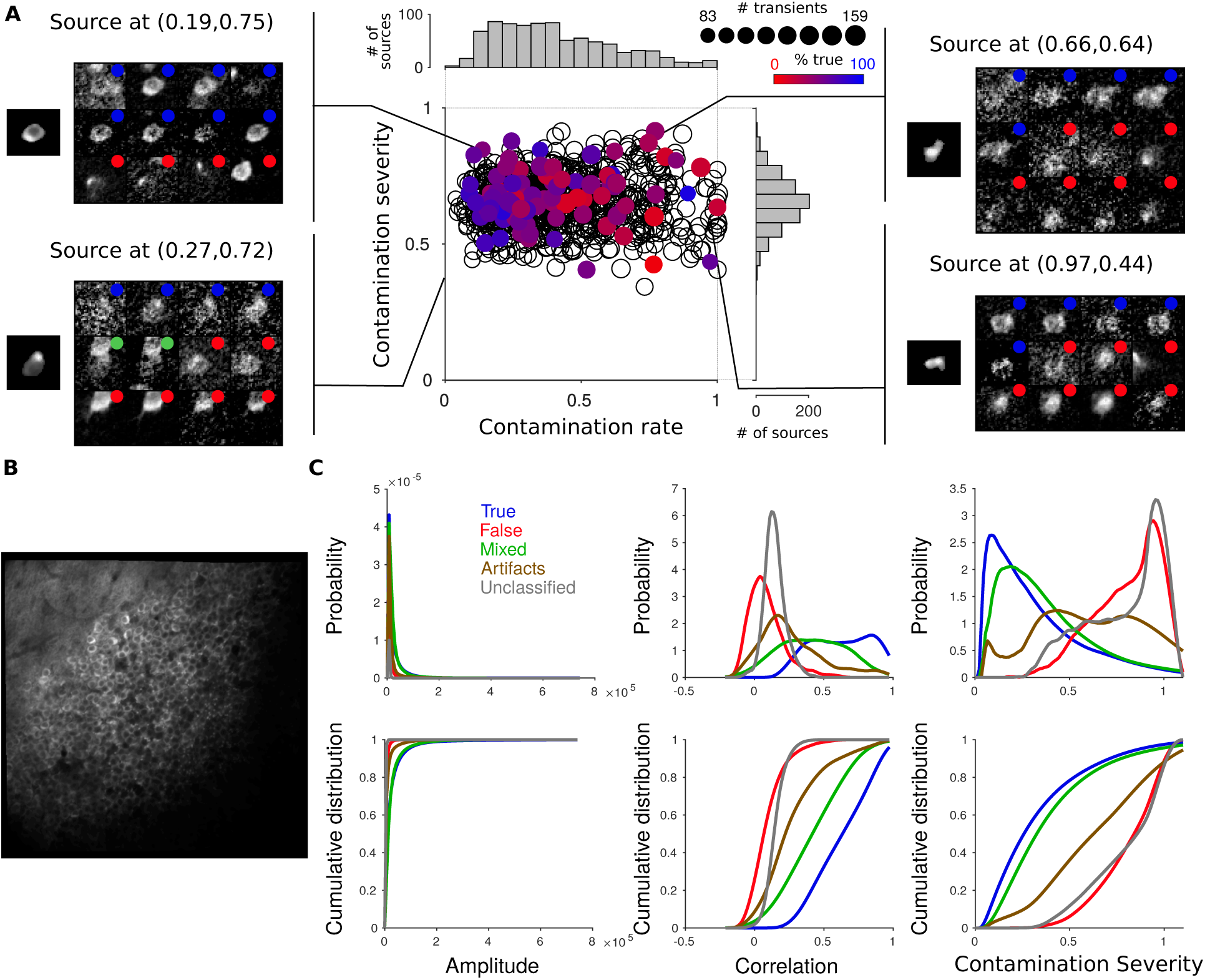
Analysis of sources estimated by CNMF from a movie of mouse CA1 densely labeled with GCaMP6f. A: FaLCon plot, same conventions as in Figure 2F. Black points indicate sources that were not human-classified. B: Average movie image. C: Properties of different categories of transients, with categories defined by human expert classification.

**Supplementary Figure 10:**
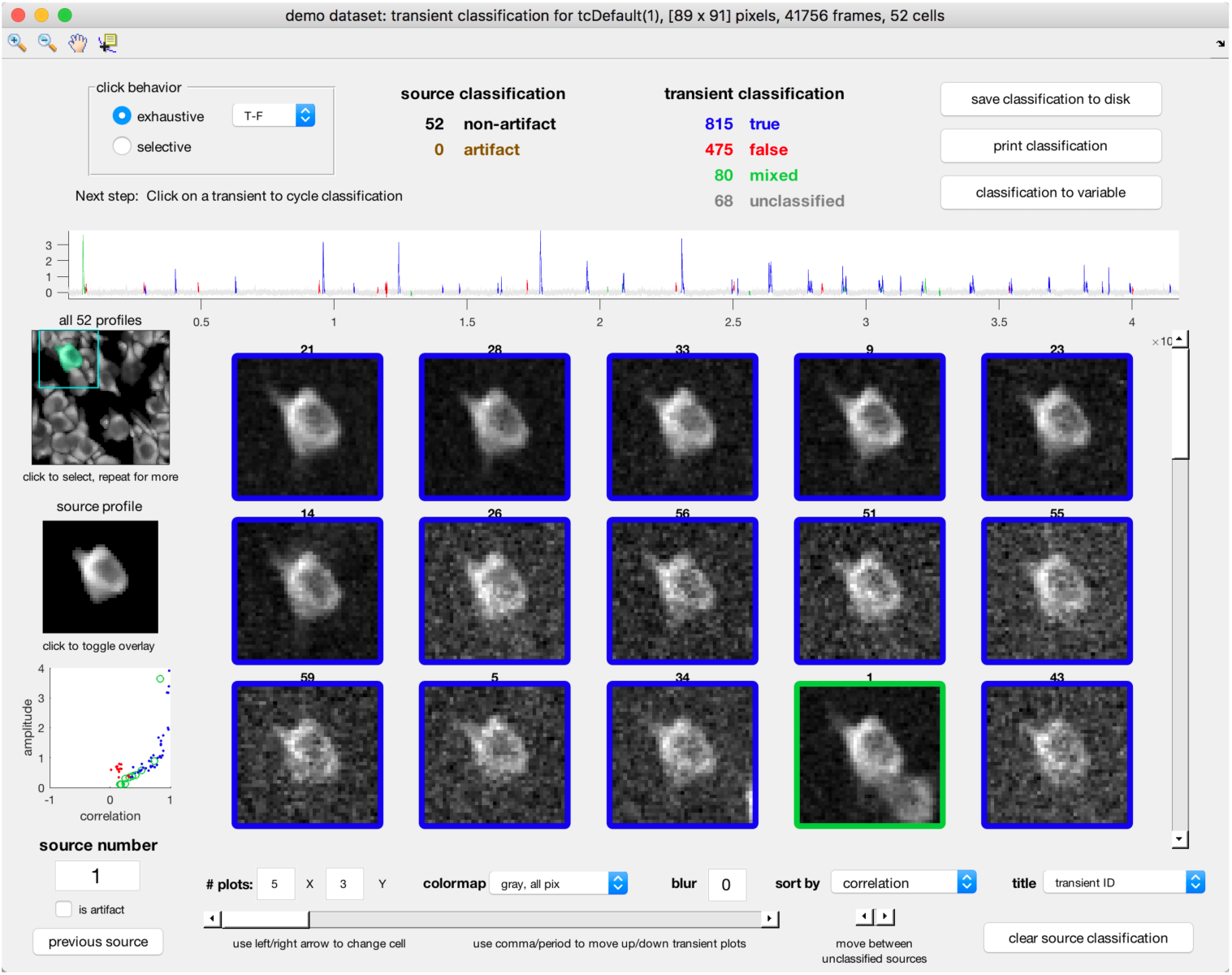
User interface for manual classification of transients.

**Supplementary Figure 11:**
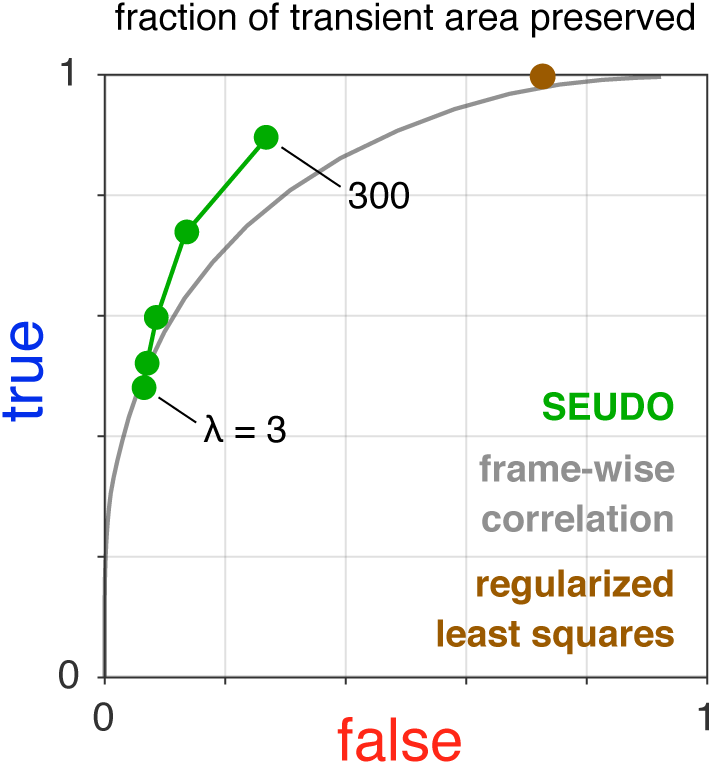
Performance of SEUDO compared to simple frame-wise classification algorithm, as measured by integrating total transient area. Same sources and transients as in Figure 6C.

**Supplementary Figure 12:**
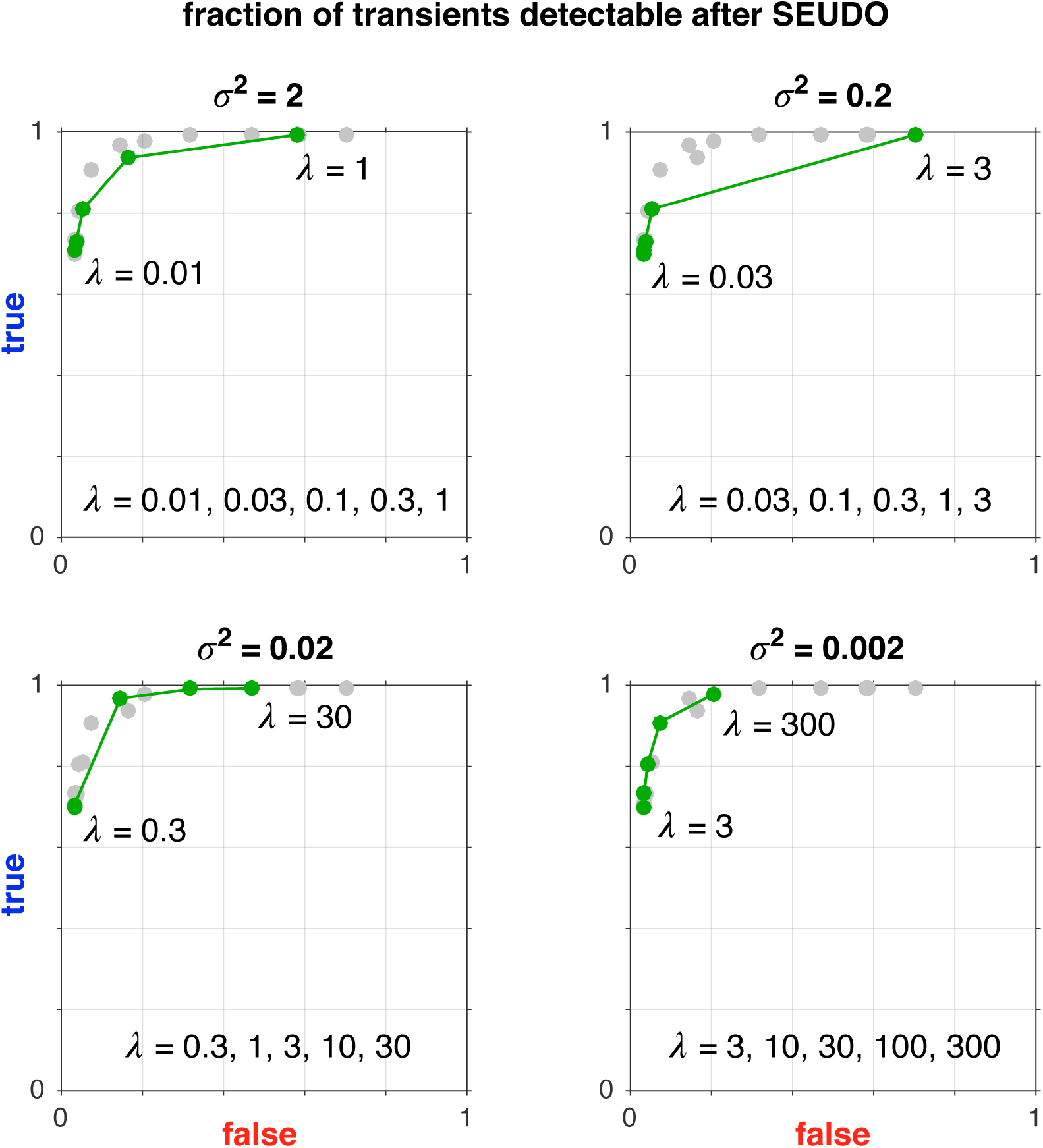
Performance of SEUDO as σ^2^ was varied over three orders of magnitude, for the same sources and quantified in the same way as in Figure 6C. Each subplot shows performance for the value of σ^2^ indicated in the title and the shown values of *λ* (green points). Also plotted for comparison are the collection of points taken from all subplots (gray points). In each case, altering *λ* biased filtering toward either preserving true transients or rejecting false transients. Each value of σ^2^ yielded approximately similar separation of true and false transients, as indicated by overlap of green and gray points, though in some cases the biasing shifted more rapidly as *λ* was varied (such as σ^2^ = 0.2).

**Table 1:** Glossary of terms

| **Term** | **Definition** |
| --- | --- |
| source | cell body, dendrite, axon, neuropil, or any other active component; this can also refer to the *estimate* of a source produced by a cell finding algorithm |
| source time course | a source's overall fluorescence value at each time point |
| source profile | spatial weight of a source in each pixel |
| transient | a discrete event, defined by the timespan in which the source time course is elevated above baseline; if sufficiently frequent, single transients might not be distinguishable |
| transient time course | the source time course during a specific transient |
| transient profile | weighted average of frames during a transient |
| true transient | transient that arose from the source it was assigned to |
| false transient | transient that did not arise from the source it was assigned to |
| mixed transient | combination of a true and false transient overlapping in time |
| fused source | source whose profile is the combination of what should be two or more distinct sources |
| artifact source | source produced by an artifact, e.g., a motion artifact, or a fused source |

## References

[1] Aleksandra Badura, Xiaonan R Sun, Andrea Giovannucci, Laura A Lynch, and Samuel S H Wang. Fast calcium sensor proteins for monitoring neural activity. Neurophotonics, 1(2):025008, 2014.

[2] Tsai-Wen Chen, Trevor J Wardill, Yi Sun, Stefan R Pulver, Sabine L Renninger, Amy Baohan, Eric R Schreiter, Rex A Kerr, Michael B Orger, Vivek Jayaraman, et al. Ultrasensitive fluorescent proteins for imaging neuronal activity. Nature, 499(7458):295, 2013.

[3] Marlene R Cohen and Adam Kohn. Measuring and interpreting neuronal correlations. Nature neuroscience, 14(7):811, 2011.

[4] H. Dana, T.-W. Chen, A. Hu, B. C Shields, C. Guo, L. L. Looger, D. S. Kim, and K. Svoboda. Thy1-gcamp6 transgenic mice for neuronal population imaging in vivo. PLoS One, 9(9):e108697, 2014.

[5] Winfried Denk, James H Strickler, and Watt W Webb. Two-photon laser scanning fluorescence microscopy. Science, 248(4951):73–76, 1990.

[6] D. A. Dombeck, C. D. Harvey, L. Tian, L. L. Looger, and D. W. Tank. Functional imaging of hippocampal place cells at cellular resolution during virtual navigation. Nature neuroscience, 13(11):1433–1440, 2010.

[7] Daniel A Dombeck, Michael S Graziano, and David W Tank. Functional clustering of neurons in motor cortex determined by cellular resolution imaging in awake behaving mice. Journal of Neuroscience, 29(44):13751–13760, 2009.

[8] Daniel A Dombeck, Anton N Khabbaz, Forrest Collman, Thomas L Adelman, and David W Tank. Imaging large-scale neural activity with cellular resolution in awake, mobile mice. Neuron, 56(1):43–57, 2007.

[9] Alexander S Ecker, Philipp Berens, Georgios A Keliris, Matthias Bethge, Nikos K Logothetis, and Andreas S Tolias. Decorrelated neuronal firing in cortical microcircuits. science, 327(5965):584–587, 2010.

[10] Jeffrey L Gauthier and David W Tank. A dedicated population for reward coding in the hippocampus. Neuron, 99(1):179–193, 2018.

[11] Andrea Giovannucci, Johannes Friedrich, Pat Gunn, Jeremie Kalfon, Sue Ann Koay, Jiannis Taxidis, Farzaneh Najafi, Jeffrey L Gauthier, Pengcheng Zhou, David W Tank, et al. Caiman: An open source tool for scalable calcium imaging data analysis. bioRxiv, page 339564, 2018.

[12] Kenneth D Harris, Rodrigo Quian Quiroga, Jeremy Freeman, and Spencer L Smith. Improving data quality in neuronal population recordings. Nature neuroscience, 19(9):1165, 2016.

[13] Espen J Henriksen, Laura L Colgin, Carol A Barnes, Menno P Witter, May-Britt Moser, and Edvard I Moser. Spatial representation along the proximodistal axis of ca1. Neuron, 68(1):127–137, 2010.

[14] Daniel N Hill, Samar B Mehta, and David Kleinfeld. Quality metrics to accompany spike sorting of extracellular signals. Journal of Neuroscience, 31(24):8699–8705, 2011.

[15] Hakan Inan, Murat A Erdogdu, and Mark Schnitzer. Robust estimation of neural signals in calcium imaging. In Advances in Neural Information Processing Systems, pages 2905–2914, 2017.

[16] Sander W Keemink, Scott C Lowe, Janelle MP Pakan, Evelyn Dylda, Mark CW van Rossum, and Nathalie L Rochefort. Fissa: A neuropil decontamination toolbox for calcium imaging signals. Scientific reports, 8(1):3493, 2018.

[17] Aaron M Kerlin, Mark L Andermann, Vladimir K Berezovskii, and R Clay Reid. Broadly tuned response properties of diverse inhibitory neuron subtypes in mouse visual cortex. Neuron, 67(5):858–871, 2010.

[18] Jason ND Kerr, David Greenberg, and Fritjof Helmchen. Imaging input and output of neocortical networks in vivo. Proceedings of the National Academy of Sciences, 102(39):14063–14068, 2005.

[19] Jérôme Lecoq, Joan Savall, Dejan Vucinic, Benjamin F Grewe, Hyun Kim, Jin Zhong Li, Lacey J Kitch, and Mark J Schnitzer. Visualizing mammalian brain area interactions by dual-axis two-photon calcium imaging. Nature neuroscience, 17(12):1825, 2014.

[20] L. Madisen, A. R. Garner, D. Shimaoka, A. S. Chuong, N. C. Klapoetke, L. Li, A. van der Bourg, Y. Niino, L. Egolf, C. Monetti, H. Gu, M. Mills, A. Cheng, B. Tasic, T. N. Nguyen, S. M. Sunkin, A. Benucci, A. Nagy, A. Miyawaki, F. Helmchen, R. M. Empson, T. Knopfel, E. S. Boyden, R. C. Reid, M. Carandini, and H. Zeng. Transgenic mice for intersectional targeting of neural sensors and effectors with high specificity and performance. Neuron, 85(5):942–58, 2015.

[21] Wasim Q Malik, James Schummers, Mriganka Sur, and Emery N Brown. Denoising two-photon calcium imaging data. PloS one, 6(6):e20490, 2011.

[22] Ruben Martinez-Cantin. Bayesopt: A bayesian optimization library for nonlinear optimization, experimental design and bandits. The Journal of Machine Learning Research, 15(1):3735–3739, 2014.

[23] Michael B McCoy and Joel A Tropp. Sharp recovery bounds for convex demixing, with applications. Foundations of Computational Mathematics, 14(3):503–567, 2014.

[24] Gal Mishne, Ronald R Coifman, Maria Lavzin, and Jackie Schiller. Automated cellular structure extraction in biological images with applications to calcium imaging data. bioRxiv, page 313981, 2018.

[25] Eran A Mukamel, Axel Nimmerjahn, and Mark J Schnitzer. Automated analysis of cellular signals from large-scale calcium imaging data. Neuron, 63(6):747–760, 2009.

[26] Junichi Nakai, Masamichi Ohkura, and Keiji Imoto. A high signal-to-noise ca 2+ probe composed of a single green fluorescent protein. Nature biotechnology, 19(2):137, 2001.

[27] John O’Keefe. Place units in the hippocampus of the freely moving rat. Exp. Neurol., 51:78—-109, 1976.

[28] Marius Pachitariu, Carsen Stringer, Sylvia Schröder, Mario Dipoppa, L Federico Rossi, Matteo Carandini, and Kenneth D Harris. Suite2p: beyond 10,000 neurons with standard two-photon microscopy. Biorxiv, page 061507, 2016.

[29] Simon P Peron, Jeremy Freeman, Vijay Iyer, Caiying Guo, and Karel Svoboda. A cellular resolution map of barrel cortex activity during tactile behavior. Neuron, 86(3):783–799, 2015.

[30] Ashley Petersen, Noah Simon, and Daniela Witten. Scalpel: Extracting neurons from calcium imaging data. arXiv preprint arXiv:1703.06946, 2017.

[31] E. A. Pnevmatikakis, D. Soudry, and et al. Simultaneous denoising, deconvolution, and demixing of calcium imaging data. Neuron, 89(2):285–299, 2016.

[32] Juan Prada, Manju Sasi, Corinna Martin, Sibylle Jablonka, Thomas Dandekar, and Robert Blum. An open source tool for automatic spatiotemporal assessment of calcium transients and local ‘signal-close-to-noise’activity in calcium imaging data. PLoS computational biology, 14(3):e1006054, 2018.

[33] John Peter Rickgauer, Karl Deisseroth, and David W Tank. Simultaneous cellular-resolution optical perturbation and imaging of place cell firing fields. Nature neuroscience, 17(12):1816, 2014.

[34] Robert Tibshirani. Regression shrinkage and selection via the lasso. Journal of the Royal Statistical Society. Series B (Methodological), pages 267–288, 1996.

[35] Photomultiplier Tubes. Basics and applications. Hamamatsu Photonics KK, Iwata City, pages 438–0193, 2006.

[36] Deborah S Won, David Y Chong, and Patrick D Wolf. Effects of spike sorting error on information content in multi-neuron recordings. In Neural Engineering, 2003. Conference Proceedings. First International IEEE EMBS Conference on, pages 618–621. IEEE, 2003.

[37] Rafael Yuste and Winfried Denk. Dendritic spines as basic functional units of neuronal integration. Nature, 375(6533):682, 1995.

